# Zinc-finger proteins with a co-opted capsid domain anchor nucleosomes over transposon sequences

**DOI:** 10.1101/2025.03.03.638093

**Authors:** Wayo Matsushima, Julien Duc, Shaoline Sheppard, Cyril Pulver, Delphine Grun, Sandra Offner, Charlène Raclot, Evarist Planet, Didier Trono

## Abstract

Transposable elements (TEs) are frequently co-opted as *cis*-regulatory sequences that govern multiple aspects of host biology. The regulatory activity of these domesticated sequences are controlled by host factors, notably KRAB domain-containing zinc-finger proteins (KZFPs) in tetrapods. Here, we report that SCAN domain-containing zinc-finger proteins (SZFPs), which originally arose through capture of a retroviral capsid domain by a KZFP gene, have expanded and diversified their DNA recognition specificity to bind distinct TE subfamilies. We further demonstrate that SZFPs anchor nucleosomes at their target sites, and that their depletion leads to global shifts of nucleosomes away from underlying TE-derived sequences, occasionally accompanied by a gain of enhancer-associated chromatin states. Thus, SZFPs represent a novel layer of chromatin regulation centered on rapidly evolving TE-derived regulatory sequences.

## Introduction

Throughout evolution, transposable elements (TEs) have been co-opted (*1*) to provide either novel genes, parts thereof (*2*), or, more often, *cis*-regulatory sequences that exert key influences on host biology (*3*). Owing to their lineage-specific emergence and expansion, TEs contribute to the formation of species-specific gene regulatory networks. Up to 25% of human candidate *cis*-regulatory elements are derived from TEs, and more than 90% of these are lineage- or even species-specific (*4*). The uncontrolled activity of TE-derived regulatory sequences is highly disruptive for host biology from the earliest stages of embryogenesis (*5*), so their domestication requires *trans*-acting modulators.

KRAB domain-containing zinc-finger proteins (KZFPs) represent a paradigmatic example of factors controlling the regulatory potential of TEs (*6*). KZFP genes first emerged in the common ancestor of *Sarcopterygii* and subsequently underwent lineage-specific expansions to generate ∼400 paralogues in the human genome alone (*7*). Most human KZFPs recognise a specific TE subfamily as a primary target (*7, 8*). Some 90% act as transcriptional repressors through the KRAB-mediated recruitment of TRIM28 (also known as KAP1) (*9, 10*), which nucleates a H3K9me3 heterochromatin-inducing complex comprising notably SETDB1 (*11*) and HP1 (*12, 13*). The vast majority of KZFPs encoded in the human genome target TE-derived sequences that are highly degenerate and incapable of transposition (*7, 8, 14*). Thus, most KZFP genes have likely been fixed in our genome independently of transposition control. Illustrating this hypothesis, a mouse KZFP gene, *Zfp568*, was found to be essential for the control of a single host gene, *Igf2*, without any significant impact on TE expression (*15*). We and other groups have also identified a number of instances where KZFPs generally influence human gene expression by targeting TE-derived regulatory sequences in multiple contexts from early embryogenesis to neuronal function (*16*–*23*). Thus, the continuous expansion and evolution of KZFPs to target rapidly evolving TEs may have facilitated the co-option of TE-derived elements, thereby contributing to the evolution of lineage-specific regulatory networks while creating the ground for selection of new phenotypic traits.

While KZFPs represent a primary example, such co-evolution with TE might not be unique to this gene family. A strong positive correlation between the numbers of LTR elements and host tandem ZF genes was observed across vertebrates including species completely lacking KZFP genes (*24*). In line with this, a recent study discovered that a large zinc-finger proteins (ZFP) family marked with another domain, named FiNZ, binds and controls TEs in cyprinid fish (*25*), suggesting that TE-controlling gene families are widespread in higher organisms. Notably, an ancestral KZFP gene gave rise to another group of ZFPs through the capture of a sequence encoding the C-terminal domain of the capsid protein of an endogenous retrovirus, Gmr1-like, as the SCAN domain (*26*). This resulted in the ancestral gene encoding a SCAN domain-containing zinc-finger protein (SZFP), and it subsequently experienced multiple rounds of duplication with frequent losses of the KRAB domain, forming a gene family widely distributed in amniotes, including 55 representatives in the human genome. Individual SZFP members have been investigated and found to play crucial roles in various biological processes, including myelination, genomic imprinting, neural development, spermatogenesis, telomere elongation, and nucleolar targeting (*27*–*32*). However, neither the evolutionary trajectory that led to such a large gene family nor any unifying function for its products has been unraveled to date.

Here, we comprehensively characterised SZFPs as a family using a combination of comparative genomic and biochemical approaches. We discovered that this transcription factor (TF) family has expanded and diversified through targeting TE-derived elements as observed for KZFPs. We also found that SZFPs anchor neighbouring nucleosomes over their target TE-derived sequences and thereby prevent them from gaining enhancer-associated chromatin states without introducing major repressive chromatin marks including H3K9me3 and H3K27me3. Our work thus identifies SZFPs as a novel class of transcriptional controllers aimed at TE-derived regulatory sequences contributing to the rapid and lineage-specific evolution of gene regulatory networks.

## Results

### SCAN-encoding elements have expanded in a lineage-specific fashion

To delineate the emergence and expansion of the SCAN domain-encoding elements, we profiled the corresponding sequences across 68 vertebrate genomes. Since SCAN domains could be a part of either the original LTR retrotransposons, Grm1-like, or ZFPs (*26*), we systematically examined their downstream regions. Gmr1-like is characterised by its unique domain organisation, distinct from that of its closest relative, Ty3/mdg4 (formerly known as Ty3/gypsy) (*33, 34*). We correspondingly found SCAN-coding sequences followed by protease (PR), integrase (IN), and RT/RNase H (RT-RH) retroviral domains (fig. S1A), which matched the previously reported domain organisation of Gmr1-like, and also cases where they were upstream of ZFP-coding sequences with or without intervening KRAB domain (Fig. 1A). This analysis revealed a striking contrast between Gmr1-like and SZFPs (Fig. 1B). Gmr1-like elements were particularly abundant in fish and turtle genomes but depleted in mammalian and reptilian genomes, suggestive of independent losses. Conversely, SZFPs were mostly present in amniotes with a marked expansion in squamate genomes. As previously observed for KZFPs (*7*), birds have lost the SZFP gene family almost entirely, with only one paralogue remaining.

**Figure 1:**
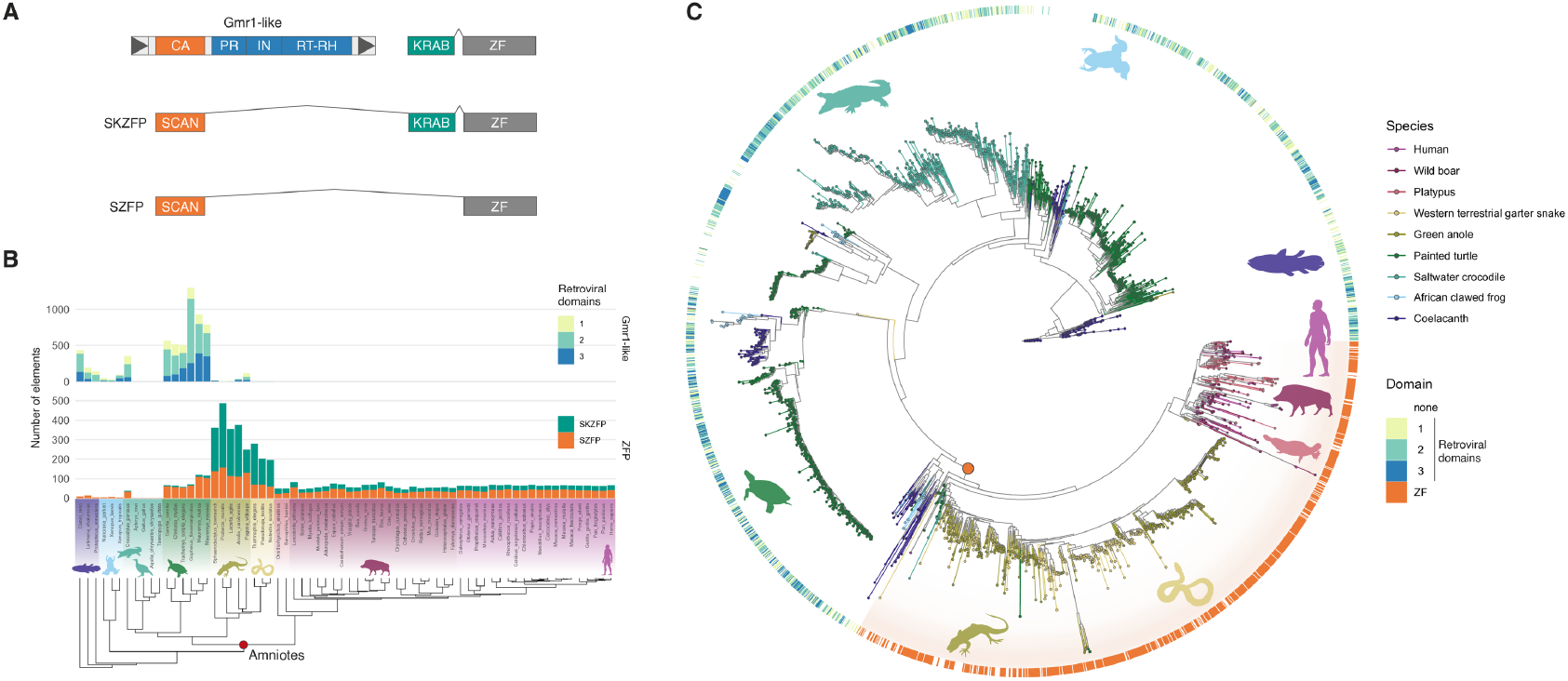
Vertebrate-wide expansion of SCAN-encoding elements. (A) Schematic of three major forms of SCAN-encoding elements: Gmr1-like, ZFP with both SCAN and KRAB domains (SKZFP), and SCAN-only ZFP (SZFP). This also represents the evolutionary trajectory of these from top to bottom. (B) The numbers of SCAN-encoding elements across vertebrate genomes. (C) Phylogenetic tree of SCAN domains from representative vertebrate species. The outer ring indicates associated domains. The orange point represents a clade including all the ZF-associated SCAN domains and is supported with SH-aLRT = 98.4%.

We then constructed a phylogenetic tree of SCAN domain sequences from representative vertebrate species to trace their evolutionary trajectory across these lineages (Fig. 1C). SCAN domains not part of Gmr1-like in amniotes formed a distinct monophyletic tree (SH-like approximate likelihood ratio test (SH-aLRT) = 98.4%). Since this clade includes SCANs either associated with a zinc-finger (ZF) array or encoded by non-ZF-associated host genes (e.g. *SCAND1*), it suggests that the acquisition of SCAN as a host protein domain has occurred only once, as previously proposed based on a limited number of species (*26*). Further, the phylogenetic tree shows that independent SZFP gene duplications took place in mammals and squamates, with specific expansions also occurring in snakes after their separation from lizards. These data indicate rapid lineage-specific expansions of SZFP genes.

We also compared the consensus sequences of the Gmr1-like capsid and the SCAN domain and observed multiple residues overrepresented in the co-opted domains compared to their ancestral retroviral counterparts (fig. S1B). Although some of these may be due to a selective sweep, the SCAN K41 residue, which displayed strong enrichment in SZFP genes, has been reported to be involved in SCAN dimerisation (*35, 36*) and has been under one of the strongest purifying selections (further discussed below). Thus, some of these amino-acid changes may have facilitated its domestication as a ZF-associated domain.

### The majority of SZFPs do not interact with TRIM28

We next turned our attention to the 58 SCAN domain-encoding genes in the human genome. We found their majority to be organized in gene clusters on chromosomes 3, 6, 7, 16, 18, and 19, suggesting expansion by tandem duplications (fig. S2A). Consistent with the previous finding that the single ancestral SZFP gene emerged as the SCAN-KRAB-ZF domain order configuration through the capture of an upstream Gmr1-like element by a KZFP gene (*26*), all SZFPs (55 genes) carry the ZF array at their C-terminus with a KRAB situated in between for the 24 proteins that still harbour this domain (fig. S2B, C).

We have previously reported that the majority of the KRAB domains present in human SZFPs are highly divergent from the consensus and are unable to interact with TRIM28 (*9*). This observation was validated by a structural study of the KRAB-TRIM28 interaction, which confirmed that many KRAB domains in SZFPs contain multiple mutations in residues critical for the KRAB-TRIM28 interaction (*37*). To comprehensively classify human SZFPs based on their ability to recruit TRIM28, we combined multiple sources of information, including KRAB amino-acid sequences, two reporter-based repression assay datasets (*10, 38*), and two protein interactome datasets (*9, 39*). We concluded that only seven SZFPs, ZFP69B, ZNF197, ZNF215, ZNF274, ZNF287, ZNF445, and ZNF75D, interact with TRIM28 with high confidence (table S1). The remaining 17 other KRAB-containing SZFPs and 31 KRAB-less SZFPs are not TRIM28-interacting, suggesting that the vast majority of SZFPs have expanded and been fixed in the genome independently of TRIM28 recruitment. This warrants a separate investigation of SZFPs from KZFPs, most of which possess a TRIM28-recruiting KRAB domain (*10*).

### SCAN is a co-opted dimerisation domain under purifying selection

The protein interactome data suggested that, unlike KZFPs, there was no single protein interactor shared by the majority of SZFPs (fig. S3A). Instead, each SZFP interacted with multiple SCAN-containing proteins with high confidence, and the protein domain most frequently identified in SZFP preys was the SCAN domain (fig. S3B). Consistent with previous studies reporting that the SCAN-SCAN homo-dimerisation is selective (*35, 40, 41*), some combinations of SZFPs appeared to associate more stably (fig. S3C).

To put the above observations into an evolutionary perspective, we calculated the ratio of non-synonymous to synonymous substitutions (*d*N/*d*S) for each amino-acid residue of SZFPs found across the 68 species used in our phylogenetic study. We expectedly detected strong signs of purifying selection for the KRAB domains of TRIM28-interacting SZFPs (fig. S4A), whereas it was the case for only a few of those not interacting with TRIM28 (fig. S4A-C). It suggests that the latter KRAB domains are largely non-functional with only a minority recently neo-functionalised as observed for some KZFPs (*9, 10*). By contrast, the vast majority of SCAN domains were under strong purifying selection regardless of the presence of an adjacent KRAB domain (fig. S4A, D). Within SCAN-encoding sequences, significantly stronger purifying selection was observed at residues involved in the dimerisation interface (*35, 36*) (fig. S4E, F). Thus, the SCAN domain might have been fixed in the host genome to facilitate dimerisation of SZFP proteins.

### SCAN is associated with lineage-specific zinc-finger repertoires

We next characterised the ZF array associated with SZFPs. The number of ZF domains in human SZFPs is on average lower than KZFPs, but still significantly higher than other ZFPs (Fig. 2A). Previous structural analyses (*42, 43*) and biochemical assays (*44*) have determined that the residues at the −1, 2, 3 and 6 positions of the alpha helix are responsible for base recognition specificity of a Cys2-His2 (C2H2) ZF domain. Thus, the similarity of the target sequences of two ZFPs can be assessed by comparing sets of the DNA-interacting residues in their ZF arrays (hereafter ZFprint) (*7, 45*) (fig. S5A). We performed this pairwise ZF identity analysis, comparing all the human ZFPs with those encoded in 64 other vertebrate genomes (fig. S5B). We attributed ages to human ZFPs based on the most evolutionarily distant species in which a ZFP with an identical ZFprint was found. Strikingly, while the majority of ZFPs without KRAB or SCAN domains were endowed with highly conserved ZFprints amongst all examined vertebrates, those found in the human KZFPs and SZFPs were largely restricted to mammals, and a number of them were primate- or human-specific (Fig. 2B, fig. S5B). This indicates that SZFPs have experienced a similar rapid turnover as KZFPs, in sharp contrast with other ZFPs, which are generally highly conserved.

**Figure 2:**
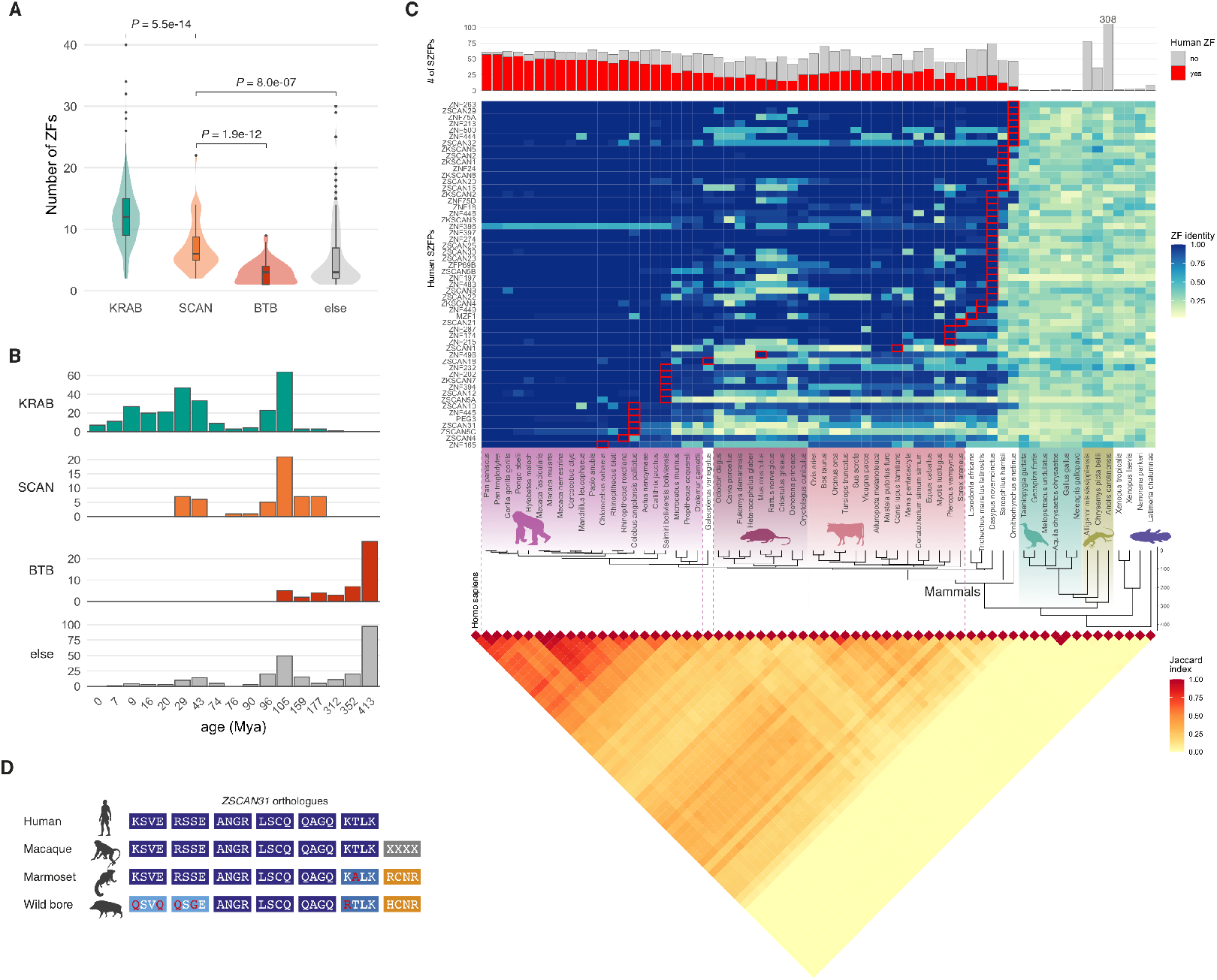
Evolutionary trajectory of the DNA-contacting residues of the human SZFPs. (A) Violin plot of the number of ZF domains associated with different protein domains encoded in the human genome. *P* values obtained from a Mann-Whitney *U* test are shown. (B) ZF age histogram of the human ZFPs, based on the last common ancestor with the most distant species harbouring the identical ZFprint. ZFPs bearing both the KRAB and SCAN domains are included in the SCAN group. (C) The numbers of SZFP genes encoded in vertebrate genomes. Of those, ZFPs bearing a ZFprint identical to the human SZFPs are highlighted in red (top). Heatmap summarising the ZF identity of the human SZFPs compared across ZFPs encoded in other vertebrate genomes. Tiles with red borders highlight the most evolutionarily distant species that harbours a ZFP with the identical ZFprint (middle). Heatmap summarising the Jaccard index calculated by pairwise comparisons of ZFprint repertoires (bottom). (D) ZFprint encoded by *ZSCAN31* orthologues from four representative species.

Among human SZFPs, we found their ZFprints conserved across placental mammals in more than half of the cases, but little homology to reptilian ZFPs, despite the high abundance of SZFP genes in this clade (Fig. 2C). This is consistent with major and independent SZFP duplications in the mammalian and reptilian common ancestors revealed by the sequence analysis of the SCAN domains shown above (Fig. 1C). In addition, we discovered more recently emerged SZFP genes in the human genome; for instance, *ZSCAN5A* and *ZSCAN5C* are the simian-restricted paralogues of *ZSCAN5B*, further refining their previous definition as primate-specific (*46*). In between these two extreme groups of SZFPs, there are also a number of evolutionarily old SZFPs that have changed their ZFs in a lineage-specific fashion. *ZSCAN31*, for example, has repetitively evolved its ZFprints since its emergence in the common ancestor of placental mammals (Fig. 2D), suggesting continuous positive selection. Overall, the repertoire of SZFP ZFprints across vertebrate genomes is highly diverse, with only a small fraction shared with species from different clades (Fig. 2C, bottom). Thus, while the majority of the human SZFP genes are conserved across placental mammals, some display signs of continuous evolution, suggesting that their targets are also rapidly evolving.

### SZFPs bind nucleosome-associated TE subfamilies of matching evolutionary ages

To identify the genomic targets of all human SZFPs, we established 55 293T cell lines, each overexpressing one of these proteins tagged with a haemagglutinin (HA) epitope, and subjected these cells to chromatin immunoprecipitation followed by sequencing (ChIP-seq). To achieve stringent identification of SZFP binding sites, we only retained samples in which a motif significantly correlated with its prediction based on the ZFprint (*47*). We also applied the same pipeline on publicly available ENCODE ChIP-seq datasets (*48*) and used it instead when a motif with a higher AUC score was obtained (table S2). Despite independent experiments performed under different conditions, the motifs obtained for the same protein were overall highly correlated between the datasets (fig. S6A). The analysis revealed that, although SZFPs are all paralogous to each other, their DNA target motifs are largely dissimilar (MoSBAT score < 0.5) even for proteins encoded within a same gene cluster, with the exception of the *ZNF75A-ZNF75D* paralogue pair (fig. S6B).

We then examined the genome-wide binding patterns of the human SZFPs. The genomic features bound by SZFPs were highly heterogeneous; some showed a strong preference for promoter regions, while others predominantly bound intergenic regions (Fig. 3A, top). Different SZFPs rarely bound in the immediate vicinity (< 100bp) of one another, suggesting a lack of cooperative binding between paralogues (fig. S7).

**Figure 3:**
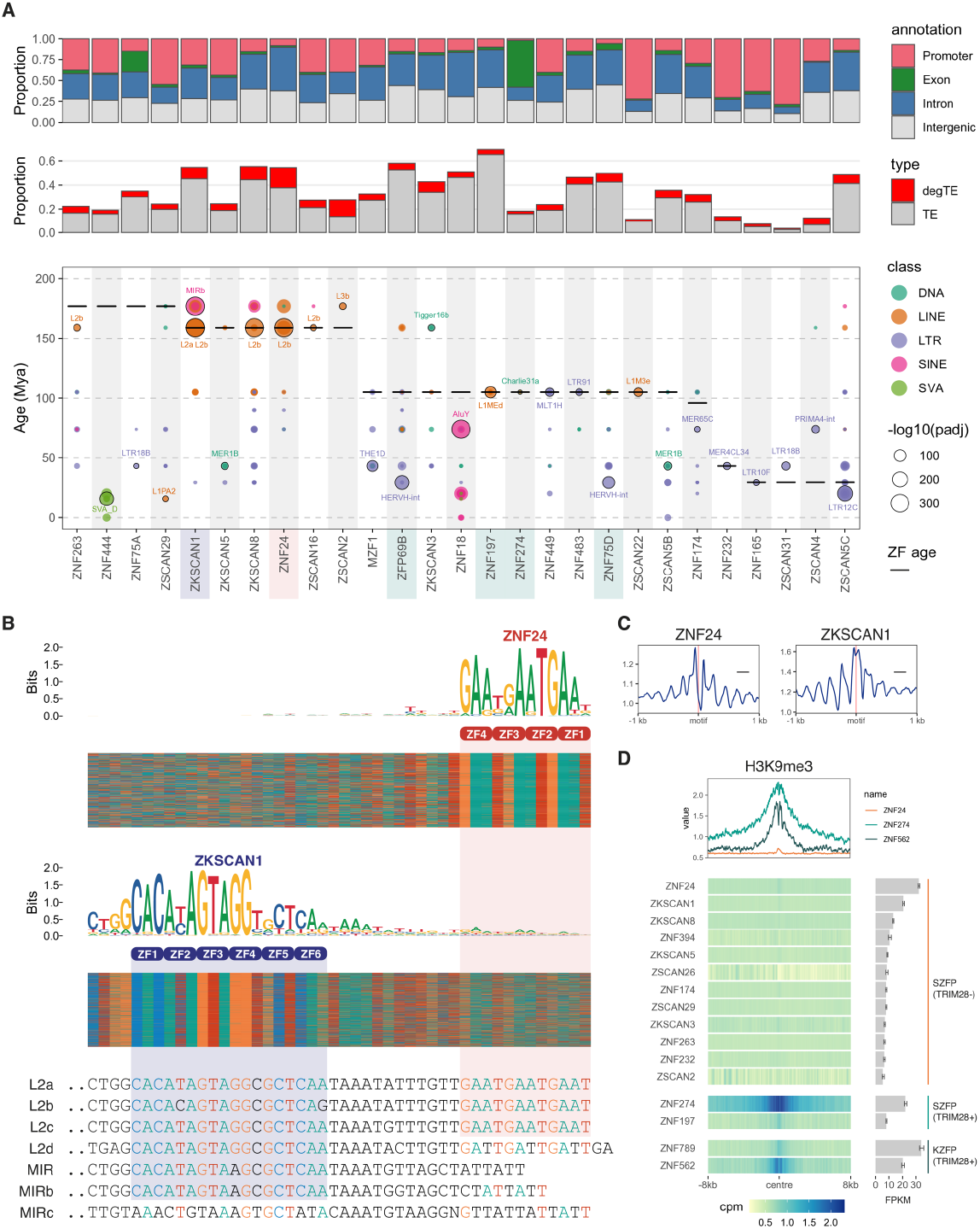
Genome-wide profiles of the human SZFP binding sites. (A) Proportions of the SZFP binding sites across different genomic regions determined by ChIP-seq. For binding sites overlapping multiple regions, priorities were given in the order shown (top). Proportions of the SZFP binding sites overlapping either TEs or degTEs (middle). TE subfamilies significantly overlap with the binding sites of the human SZFPs are shown according to their evolutionary age based on the Dfam database (bottom). Benjamini-Hochberg adjusted *P* values calculated by a binomial test using pyTEnrich are shown. TRIM28-interacting SZFPs are highlighted with a green shade. (B) TE-derived sequences bound by either ZNF24 or ZKSCAN1 are shown as heatmaps. ZF domains that were predicted to recognise a specific triplet are also shown. Consensus sequences of L2 and MIR families are shown with red and blue shades highlighting sequences recognised by ZNF24 and ZKSCAN1, respectively. (C) Nucleosome density signals around the ZNF24 and ZKSCAN1 binding motifs. (D) Heatmaps of average H3K9me3 signals (CPM) around the binding sites of SZFPs and KZFPs that are expressed in K562. The signal profiles of ZNF24, ZNF274, and ZNF562 are also shown as a line plot on the top as representatives of non-TRIM28-interacting SZFPs, TRIM28-interacting SZFPs, and KZFPs, respectively.

We next tested whether SZFPs bind TE-derived sequences (fig. S8A). We previously established a novel method to annotate highly degenerate TE-derived elements (degTEs) through computational reconstruction of ancestral genomes (*14*). The proportions of SZFP binding sites in TEs and degTEs varied significantly, ranging from ∼5% to ∼60% (Fig. 3A, middle). However, regardless of this difference, most of the examined SZFPs significantly bound at least one TE subfamily (Fig. 3A, bottom), and also the corresponding degTEs (fig. S8B). We further noted a striking correspondence between the estimated age of an SZFP and the time of emergence of its target TE subfamily (*P* = 0.03, approximative general independence test), as previously observed for KZFPs (*7*), suggesting similar modes of selection on their affinity against newly emerged TE-derived sequences. Interestingly, when several SZFPs targeted the same TE subfamily, they generally recognised distinct regions of the element. For instance, although both ZNF24 and ZKSCAN1 bound multiple L2 subfamilies, ZNF24 bound to the very 3′ end of these integrants, whereas ZKSCAN1 recognised a motif located 12 bp upstream (Fig. 3B). Accordingly, these two SZFPs targeted different sets of TE subfamilies (Fig. 3A, B). ZNF24 binding sites significantly overlapped with L2a, L2b, and L2c, but not L2d, which does not contain the (GAAT)3 target motif recognised by ZNF24. In contrast, ZKSCAN1 recognised a sequence common to all L2 subfamilies and some MIR subfamilies, which contain an L2-derived insert (*49*). Thus, the targeting of TEs by SZFPs is not only diverse but also highly specific. We also assessed the chromatin states of the SZFP binding sites using publicly available data from assay for transposase-accessible chromatin using sequencing (ATAC-seq) and ChIP-seq of histone modifications. Interestingly, SZFPs bound significantly more inaccessible genomic loci compared to other transcription factors (fig. S9A), and micrococcal nuclease sequencing (MNase-seq) data indicated regularly-spaced and phased nucleosomes around SZFP binding sites (Fig. 3C). Furthermore, unlike KZFPs, whose binding sites are enriched for H3K9me3, the binding sites of the non-TRIM28-recruiting SZFPs did not show a strong association with either this heterochromatin modification (Fig. 3D) or with H3K27me3 (fig. S9B), indicating that chromatin inaccessibility at SZFP binding sites is independent of these known repressive marks.

### SZFPs restrict chromatin accessibility through nucleosome anchoring

To investigate the functional roles of SZFP family members, we endogenously tagged *ZNF24* in human haploid HAP1 cells with an FKBP12^F36V^ degron (*ZNF24*^*FKBP*^) (fig. S10A), which allows for rapid protein degradation via recruitment to VHL by a selective small molecule, dTAG^v^-1 (*50*). We validated a ∼99 % reduction in ZNF24 protein levels after 30-min exposure to dTAG^v^-1 (fig. S10B, C). ZNF24 depletion induced chromatin remodelling around its binding sites (Fig. 4A), resulting in significant increases in distances between flanking nucleosomes (Fig. 4B). A small fraction (*n* = 105) of these loci gained chromatin accessibility as measured by ATAC-seq (Fig. 4C), including a number of L2 integrants bound by ZNF24 (Fig. 4D). The majority of these newly opened genomic loci corresponded to ZNF24 binding sites and gained H3K4me1, but rarely H3K27ac, upon depletion of this factor (Fig. 4E, F). It suggests that these loci largely became primed but not active enhancers, consistent with the observed minimal changes in gene expression (fig. S11A, B). Of note, the first intron of one of two significantly downregulated genes, *PCGF3*, was bound by ZNF24 and significantly gained accessibility, suggesting that this transcriptional change might be a direct consequence of ZNF24 depletion (fig. S11C). To examine the kinetics of the chromatin opening upon the loss of ZNF24, we performed a time course experiment (fig. S12A). Strikingly, chromatin accessibility increased after only 2 hours of exposure to dTAG^v^-1 (Fig. 4G, fig. S12B), with some genomic loci peaking at this early time point (fig. S12C). Further, pre-treatment of *ZNF24*^*FKBP*^ cells with either a SMARCA2/4 proteolysis-targeting chimera (PROTAC), ASBI1 (*51*), or a SMARCA2/4 ATPase inhibitor, BRG1i (also known as BRM014) (*52, 53*), significantly prevented the chromatin opening upon ZNF24 depletion (Fig. 4H, fig. S13), showing that the observed increase in ATAC-seq signal reflected chromatin remodelling mediated by the SWI/SNF complex. Confirming these data, we also endogenously tagged another SZFP paralogue, *Zscan10*, with an FKBP12^F36V^ in mouse embryonic stem cells (mESCs) (fig. S14A) and similarly observed the opening of its genomic binding sites upon its degradation (fig. S14B). In this case, we detected a number of differentially expressed genes (fig. S14C), including one that harbours an intronic ZSCAN10 binding site that showed accessibility increase (fig. S14D). Therefore, blocking chromatin accessibility is not a property unique to ZNF24, but is shared with other SZFPs. Finally, to assess the relevance of ZNF24 in controlling regulatory elements in physiologically relevant contexts, we asked whether the genomic loci that became open upon ZNF24 degradation in our experimental system also can become accessible in human tissues. By analyzing the ENCODE DNase-seq dataset (*54*), we found that these loci display cell type-specific chromatin accessibility along patterns resembling the opening observed when ZNF24 was degraded in the *ZNF24*^*FKBP*^ cells (Fig. 4I). This suggests that these sequences function as regulatory elements in differentiated human tissues. Since these loci exhibit distinct openings in different cell types, other factors, such as pioneer factors, are likely involved in defining accessibility in addition to the levels of ZNF24.

**Figure 4:**
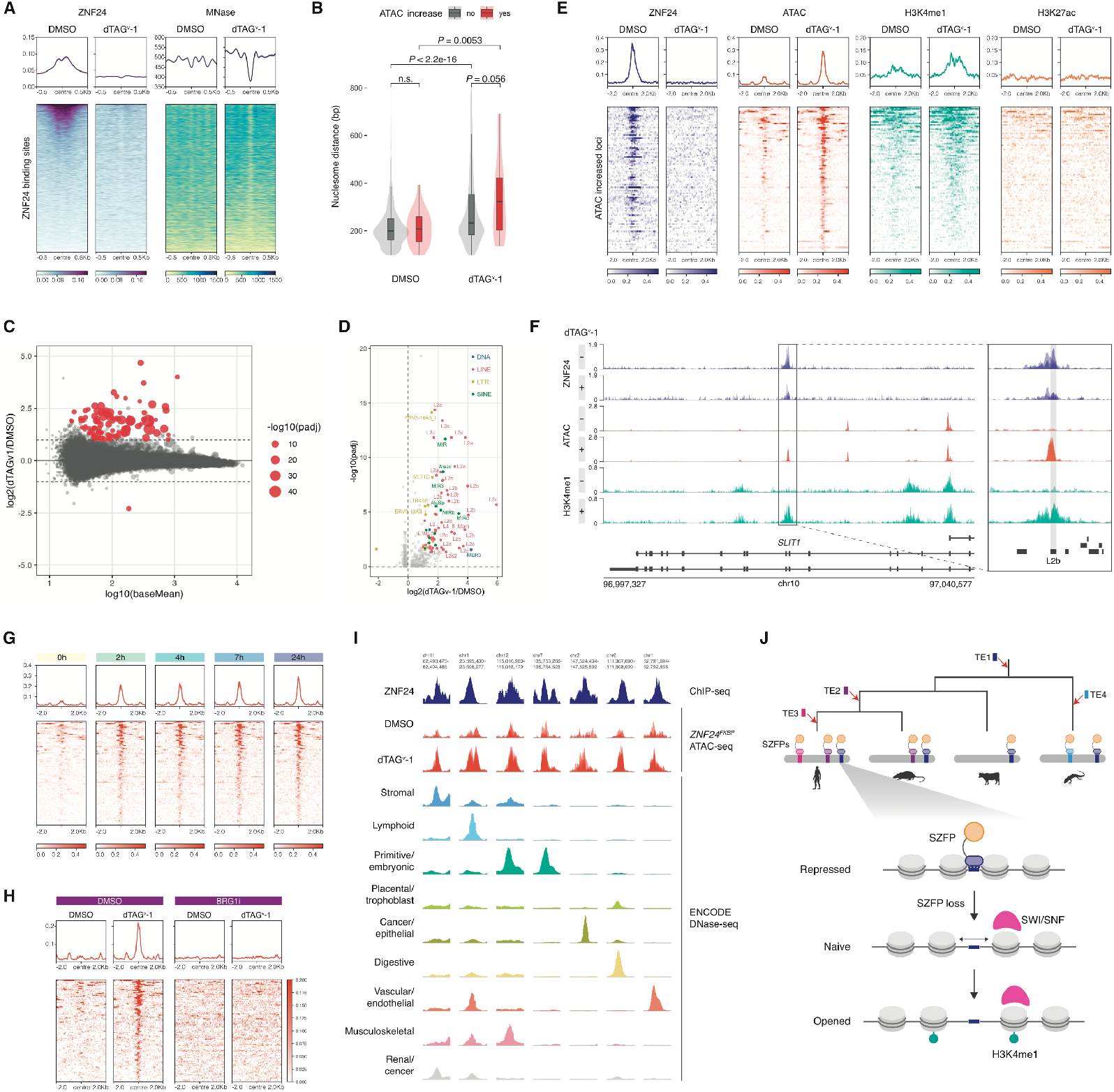
Chromatin remodelling associated with ZNF24 depletion. (A) (A) Heatmaps summarising dTAG^v^-1-induced changes of ZNF24 and nucleosomal occupancy around ZNF24 binding sites as measured by ChIPmentation and MNase-seq, respectively. (B) Violin and box plots showing nucleosome dyad distance immediately flanking ZNF24 binding sites, which are further grouped by their association with increased ATAC-seq signal. For matched and unmatched sets of genomic intervals, a Wilcoxon signed rank test and a Mann-Whitney *U* test were respectively employed for *P* value computation. (C) MA plot summarising ATAC-seq signal changes comparing *ZNF24*^*FKBP*^ HAP1 cell lines (*n* = 4) exposed either to dTAG^v^-1 or DMSO for 24 hours. Adjusted *P* values were obtained by DESeq2 with the Benajamini-Hochberg multiple testing adjustment. (D) Volcano plot summarising the changes of ATAC-seq signals in TE integrants, comparing *ZNF24*^*FKBP*^ HAP1 cell lines (*n* = 4) exposed either to dTAG^v^-1 or DMSO for 24 hours. (E) Average ChIPmentation and ATAC-seq profiles around the genomic loci where a significant accessibility gain was observed. Histone ChIPmentation was performed after 72-hour exposure to dTAG^v^-1 or DMSO. Signals were CPM normalised. (F) An example locus where a ZNF24 binding site gained accessibility upon the dTAG^v^-1 treatment. CPM normalised signals from each biological replicate are shown as transparent shades. In the zoomed window, the TE annotation is shown at the bottom. (G) Time-course of average ATAC-seq profiles around the genomic loci where a significant accessibility gain was observed. (H) Average profiles of ATAC-seq signals (CPM) around the opened regions comparing cells pre-treated with BRG1i or DMSO. (I) Example genomic loci that gain chromatin accessibility both upon the loss of ZNF24 in *ZNF24*^*FKBP*^ HAP1 cells and in specific human cells. For each genomic locus, the y-axis of each of ATAC-seq and DNase-seq signals are normalised to have the same maximum value across samples. (J) Working model for the evolution and function of SZFPs derived from this study.

Altogether, we report that SZFPs, which arose through an exonisation of a retroviral domain by a KZFP gene, have evolved to bind TEs emerging in a lineage-specific manner (Fig. 4J). We also show that SZFPs bind nucleosome-associated TE-derived sequences and prevent their target TE-derived element from gaining chromatin accessibility, independently of known heterochromatin marks. Given the extremely rapid expansion of TEs and SZFPs targeting them, SZFPs may contribute to the regulatory evolution through controlling the regulatory potential of TE-derived elements.

## Discussion

We have previously discovered that KZFPs, which constitute the largest family of transcription factors encoded by the human genome, contribute to the evolution of gene regulatory networks by partnering with their TE targets to generate a largely species-specific layer of epigenetic regulation (*7*). The present work reveals that SZFPs, an evolutionary spin-off of KZFPs, share many characteristics with KZFPs. Both families have undergone extensive lineage-specific gene expansion and rapid diversification of their ZF arrays, achieving specific targeting of TE-derived elements of a matching evolutionary age and control of regulatory potential thereof. Since we observed frequent losses or non-functionalisation of KRAB domains in SZFP genes, SZFPs may confer functional benefits, at least for some targets, over KZFPs. Since the chromatin inaccessibility induced by SZFPs does not involve H3K9me3 heterochromatinisation, which can bidirectionally spread >10 kb from an induction point (*55*), SZFPs may be able to achieve more precise control on TE-derived sequences with a minimal impact to neighbouring genetic elements. Thus, we propose that the SZFP family has also contributed to the evolution of gene regulatory networks through the targeting and control of TE-derived elements.

The mechanism of action of SZFPs provides new insights into chromatin regulation. Our data indicate that ZNF24, ZSCAN10, and a number of other SZFPs can bind nucleosome-associated DNA characterised by a lack of ATAC-seq signal. This is in line with recent *in-silico* screens, reporting several SZFPs as nucleosomal DNA binders (*56, 57*), and biochemical characterization of one family member, ZSCAN4 (*58*). TFs with nucleosomal binding capability are called “pioneer transcription factors (PTFs)”, and many of these have been shown to make chromatin accessible (*59*). Despite their nucleosomal binding capability, we observed SZFPs to have the opposite effect. In the presence of SZFPs, their targeted sequences are largely inaccessible with regularly-phased nucleosomes, and depletion of SZFPs leads to increased spacing between flanking nucleosomes, making some loci accessible in a SWI/SNF-dependent fashion. This suggests that SZFPs anchor neighbouring nucleosomes, blocking access to the underlying DNA. Although it has previously been observed that some genomic loci bearing repressive histone marks such as H3K9me3 or H3K27me3 are less frequently targeted by some PTFs (*60, 61*), we found that SZFP binding sites were not enriched for these marks, and that SZFPs did not interact with known chromatin modifiers. In addition to the two SZFPs tested in this study, another member, ZKSCAN3, was also shown to restrict chromatin accessibility of TE-derived sequences independent of its KRAB domain (*62*), suggesting that this mode of chromatin control is shared among SZFPs. Interestingly, the genomic features of SZFP binding sites resemble those recently discovered in silencer elements of the fly genome, which were found to be also recognised by polydactyl ZFPs, inaccessible by ATAC-seq, and to display phased nucleosomes around the ZFP-bound motif (*63*). Thus, SZFPs may contribute to shaping the genome-wide chromatin accessibility landscape of higher vertebrates independently of known repression mechanisms, which promises new avenues for fundamental understanding of chromatin regulation.

## Materials and Methods

### Cell culture

293T cells were cultured in Dulbecco’s modified Eagle’s medium (DMEM) (Thermo Fisher, 41966–029) supplemented with 10% fetal bovine serum (FBS) (Merck, F9665-500ML, Lot 19A124) and 1X Penicillin-Streptomycin-L-Glutamine (Corning, 30-009-CI). HAP1 cells (Horizon Discovery, C631) were cultured in Iscove’s modified Eagle’s medium (IMDM) (Thermo Fisher, 12440061) supplemented with 10% FBS (Merck, F9665-500ML, Lot #19A124) and 1X Penicillin-Streptomycin-L-Glutamine (Corning, 30-009-CI). Mouse embryonic stem cells (mESCs) were a kind gift from Professor Denis Duboule, EPFL, Switzerland and were cultured in 2i+LIF medium composed of DMEM (Thermo Fisher, 10566-024), 10% ES Cell FBS (Thermo Fisher, 16141079), 1X MEM Non-Essential Amino Acids Solution (Thermo Fisher, 11140050), 1 mM sodium pyruvate (Thermo Fisher, 11360-039), 100 µM 2-Mercaptoethanol (Thermo Fisher, 31350-010), 100 U/ml Penicillin-Streptomycin (Thermo Fisher, 15140-122), 100 ng/ml of mouse leukemia inhibitory factor (LIF) (Protein Production and Structure Core Facility, EPFL), 3 μM of GSK3 inhibitor (CHIR99021; Tocris Biosciences, 4423), and 2 μM of MEK1/2 inhibitor (PD0325901; Tocris Biosciences, 4192).

293T cells stably expressing an SZFP were generated as previously described (*7*). Gateway donor vectors encoding codon-optimised cDNA of SZFPs were obtained from GeneArt gene synthesis service (Thermo Fisher). The donor vector was shuttled using Gateway LR Clonase II enzyme mix (Thermo Fisher, 11791100) to a lentiviral transfer plasmid that expresses the insert fused with three consecutive HA tags at the C-terminal in a doxycycline-inducible manner. 1.1 µg of the resulting plasmid together with 0.7 µg R8.74, and 0.4 µg pMD2G (both plasmids are available on Addgene) were transfected to 80% confluent 293T on a 6-well plate with 6.6 µl FuGENE 6 (Promega, E2692) for lentiviral production. 48 hours after the transfection, the supernatant was recovered, filtered through a sterile 0.22-µm filter (Merck, SLGPR33RS), and added on recipient 293T cells for transduction. 72 hours after the transduction, the cells were selected with 1 µg/ml puromycin for 72 hours. Integration and expression of the SZFP transgenes were validated with Sanger sequencing and Western blot, respectively.

To establish a HAP1 cell line with its endogenous *ZNF24* gene tagged with an FKBP12^F36V^ degron, the cells were first sorted based on cellular sizes with FACSAria II (BD Biosciences) to enrich for haploid cells as previously described (*65*). The repair template plasmid construction and sgRNA cloning were performed as previously described (*66, 67*). pCRIS-PITChv2-BSD-dTAG (BRD4) was a gift from James Bradner & Behnam Nabet (Addgene plasmid # 91792 ; http://n2t.net/addgene:91792 ; RRID:Addgene_91792) and pX330-BbsI-PITCh was a gift from Peter Kaiser (Addgene plasmid # 127875 ; http://n2t.net/addgene:127875 ; RRID:Addgene_127875). The constructed pCRIS-PITChv2-BSD-dTAG (ZNF24) and pX330-BbsI-PITCh (ZNF24) were transfected to the sorted HAP1 cells with TransIT-LT1 (Mirus Bio, MIR 2306). The transfected cells were selected with 10 µg/ml blasticidin for 3 days and were clonally isolated with a serial dilution. Absence of the wild-type allele, in-frame degron insertion, and expression of tagged ZNF24 were confirmed with genomic PCR, Sanger sequencing, and Western blot, respectively.

### ChIP-seq and ChIPmentation

30 or 1 million cells were used to perform ChIP on a transcription factor and a histone modification profiling, respectively. ChIP was performed as previously described (*7*). For transcription factor overexpression, cells were cultured in 1 µg/ml doxycycline-containing culture medium for 72 hours prior to harvest. Detached cells were cross-linked with 1% (vol/vol) formaldehyde in PBS for 10 min, and the reaction was quenched with Tris-HCl (200 mM, pH = 8). The cross-linked nuclei were extracted and sonicated by the Covaris E220 Focused-ultrasonicator with the following settings: 20 min at 5% duty cycle, 140 W, 200 cycles. The fragmented chromatin was then immunoprecipitated with antibody-bound Dynabeads Protein G (Thermo Fisher, 10004D) overnight, washed with high-salt wash buffer, low-salt wash buffer, and LiCl wash buffer, and was used for either ChIP-seq or ChIPmentation library preparation as described below. Antibodies used were following: Anti-HA.11 Epitope Tag Antibody (BioLegend, 901515); H3K4me1 Antibody (Diagenode, C15410037); H3K27ac Antibody (Diagenode, C15410174).

For ChIP-seq library preparation, the chromatin was de-crosslinked with 1 µl Proteinase K (20 mg/ml) at 65°C and 1,400 rpm shaking overnight. DNA was purified with MinElute PCR purification kit (Qiagen, 28006), and 10 ng of it was used for library preparation with NEBNext Ultra End Repair/dA-Tailing Module (New England Biolabs, E7546L).

For a ChIPmentation library, the chromatin was on-bead tagmented with pA-Tn5 (Protein Production and Structure Core Facility, EPFL) on a thermomixer at 37°C for 5 min with 800 rpm shaking. The chromatin was washed twice with low-salt wash buffer and was de-crosslinked with 1 µl Proteinase K (20 mg/ml) at 65°C and 1,400 rpm shaking overnight. DNA was purified with MinElute PCR purification kit (Qiagen, 28006) and was PCR amplified with Q5 High-Fidelity 2X Master Mix (New England Biolabs, M0492L) and indexed primers. The amplified library was cleaned-up by adding 1.2 times the volume of AMPureXP beads (Beckman Coulter, A63882) and eluted with 10 mM Tris-HCl, pH 8.0.

### MNase-seq

To induce degron-tagged protein degradation, *ZNF24*^*FKBP*^ HAP1 cells were cultured in the medium containing either 500 nM dTAG^v^-1 (MedChemExpress, HY-145514) or the corresponding volume of dimethylsulfoxide (DMSO) for an indicated period of time prior to harvest. MNase-seq library preparation was performed as previously described with modifications (*68*). Briefly, nuclear extraction and permeabilisation were performed on 1.5 million detached cells in ATAC-RSB buffer containing 0.1% NP-40 and 0.1% Tween-20 on ice for 3 min. The nuclei were washed once with the same buffer and digested with 1 U MNase (Worthington Biochemical Corp., LS004798) in 50 µl buffer containing 1 mM CaCl_2_ at 37°C for 20 min on a thermomixer with 1,000 rpm mixing. The digested DNA was cleaned-up with DNA Clean and Concentrator-5 Kit (Zymo Research, D4014) and was size-selected in a 1.5% agarose gel for fragments smaller than 200 bp. The DNA was then purified from the gel, and 500 ng of it was used as input for library preparation using NEBNext® Ultra™ II DNA Library Prep Kit for Illumina (New England Biolabs, E7645S).

### ATAC-seq

To induce degron-tagged protein degradation, cells carrying an endogenous gene tagged with an FKBP12^F36V^ degron were cultured in the medium containing either 500 nM dTAG^v^-1 (MedChemExpress, HY-145514) or the corresponding volume of dimethylsulfoxide (DMSO) for an indicated period of time prior to harvest. For the SMARCA2/4 perturbation experiment, cells were exposed additionally to either 500 nM ASBI1 (MedChemExpress, HY-128359), 1 µM BRG1i (MedChemExpress, HY-119374), or corresponding volume of DMSO 24 hours prior to harvest. ATAC-seq library preparation was performed as previously described (*69*). Briefly, nuclear extraction and permeabilisation were performed on 50,000 detached cells, followed by 2.5 µl Tn5 (Illumina, 20034197) tagmentation in a 50 µl reaction at 37°C for 30 min on a thermomixer with 1,000 rpm mixing. The transposed DNA was cleaned-up with DNA Clean and Concentrator-5 Kit (Zymo Research, D4014) and was PCR amplified with Q5 High-Fidelity 2X Master Mix (New England Biolabs, M0492L) and indexed primers ordered from IDT. The amplified library was double size-selected by 0.5 and 1.5 the volumes of AMPureXP beads (Beckman Coulter, A63882) to enrich for fragments ranging from 150-700 bp and was eluted with 10 mM Tris-HCl, pH 8.0.

### RNA-seq

To induce degron-tagged protein degradation, cells carrying a gene tagged with an FKBP12^F36V^ degron were cultured in the medium containing either 500 nM dTAG^v^-1 (MedChemExpress, HY-145514) or corresponding volume of DMSO for 72 hours prior to harvest. RNA was extracted with High Pure RNA Isolation Kit (Roche, 11828665001) and 500 ng of it was used to prepare RNA-seq libraries using QuantSeq 3′ mRNA-Seq V2 Library Prep Kit (Lexogen, 191.24) following the manufacturer’s instruction.

### High-throughput DNA sequencing

All the high-throughput sequencing libraries were quantified with Qubit dsDNA Quantitation, High Sensitivity (Thermo Fisher, Q32854) and Qubit 2.0 Fluorometer (Thermo Fisher) and quality checked using High Sensitivity DNA Kit (Agilent, 5067-4626) and 2100 Bioanalyzer (Agilent). The libraries were PE75 sequenced either on a NextSeq500 (Illumina) or an AVITI (Element Biosciences) by the Gene Expression Core Facility (GECF) at EPFL. For all the runs, 1-5% PhiX was added for an increased complexity.

### Bioinformatics

#### Genome annotation

FASTA files for the selected genomes were obtained from the NCBI website (https://www.ncbi.nlm.nih.gov/). HMM profiles of SCAN (PF02023.20), zinc-finger (PF00096.29), KRAB (PF01352.30), RT_RNase (RT_RH; PF17917.4), rve (IN; PF00665.29) and RVP (PR; PF00077.23) motifs were obtained from Pfam website (http://pfam.xfam.org) and were used to identify these domains across all genomes using hmmscan function from the HMMER (http://hmmer.org/) (version 3.4) library on six-frame translated genomes as previously described (*7, 21*). Computational scripts to achieve this are available on GitHub (https://github.com/PulverCyril/kzfp_annotation). For each SCAN hit, a 10-kb downstream window was examined for presence of other previously scanned motifs to determine its association with either a Gmr1-like element or a ZFP gene. Based on the distance distribution obtained in Fig. S1A, IN and RT-RH were called associated with a SCAN domain only when they were found at a distance of < 500 bp or 600-1100 bp, respectively. A phylogenetic tree of the species used in the analysis was constructed and downloaded from the TimeTree website (*70*) (https://timetree.org/).

#### Phylogenetic analysis of SCAN sequences

SCAN domain sequences obtained as above were aligned with Clustal Omega (*71*) (version 1.2.4) with the default settings, and the alignment was used to construct a phylogenetic tree with IQ-TREE (*72*) (version 2.2.6) with the option “-alrt 1000” to perform SH-like approximate likelihood ratio test (SH-aLRT) (*73*). The resulting tree data was visualised with ggtree (*74*) (version 3.2.1).

#### ZFprint analysis

For the ZF domains annotated in the above step, DNA-contacting residues were first determined and were grouped for each ZFP gene as a ZFprint (*45*). All the human ZFprints were pairwise aligned to those in other genomes, using the pairwise2 function from Biopython (version 1.81) with a custom BLOSUM80 matrix to ensure quadruplet alignment. Computational scripts to achieve this are available on GitHub (https://github.com/PulverCyril/kzfp_annotation). A ZF identity score was calculated by dividing the number of matched DNA-binding residues by the total number of aligned residues. To estimate a time point when the human ZFprint emerged (ZF age) of a given human ZFP gene, the most distance species carrying the same ZFprint as the ZFP was determined, based on the divergence time to that species taken from the TimeTree website (*70*) (https://timetree.org/).

#### ChIP-seq analysis

Raw reads were mapped to the human genome (hg38) using Bowtie2 (*75*) (version 2.5.1), with the “--sensitive-local” mode. Before peak calling, BAM files were filtered to remove low-quality reads (MAPQ < 10) and then run through the Bioconductor package GreyListChIP (version 1.32.0) to mask regions that are known to yield artefact peaks. Peaks were called with MACS2 (*76*) (version 2.2.7.1) using the “--bampe” option, and only peaks with a score > 50 were kept for downstream analyses. The binding motifs of ZFPs were determined with RCADE2 (*47*), with an exception of ZNF444, for which STREME (*77*) from the MEME suite (version 5.5.3) was instead used. The same motif identification pipeline was applied to the ENCODE ChIP-seq peaks, and, for SZFPs with multiple data sources, the data that yielded a motif with the highest AUC was used for downstream analyses. Pairwise motif similarity analyses were performed with MoSBAT (*78*) (v1.0.0). The ChIP-seq peak sequences were searched for the presence of the corresponding SZFP motif using FIMO (*79*) from the MEME suite (version 5.5.3) with the default parameters, and the regions matching the motif within peaks were used as SZFP binding sites in the downstream analyses.

#### TE enrichment analysis

To find TE subfamilies that are significantly enriched in the binding sites of SZFPs, pyTEnrich (https://alexdray86.github.io/pyTEnrich/build/html/index.html) was used with the default parameters.

#### Epigenetic profiling of the TF binding sites

bigWig files of ATAC-seq (ENCFF102ARJ), H3K27me3 (ENCFF242ENK), and H3K9me3 (ENCFF812HRW) and an ATAC-seq peak BED file (ENCFF333TAT) in K562 were downloaded from the ENCODE website (*48*) (https://www.encodeproject.org/). ChIP-seq peak BED files of TFs were also downloaded from the same website and are listed in Data S1. SZFP expression data in K562 was downloaded from the Human Protein Atlas (*80*) (https://www.proteinatlas.org/).

#### SCAN interactome analysis

PPI in HEK293T was downloaded from the BioPlex website (*39*) (https://bioplex.hms.harvard.edu/) and was filtered for only interactions that involve SCAN-containing proteins. The resulting networks were visualised with Gephi (*81*) using the “ForceAtlas2” layout.

#### *d*_N_/*d*_S_ calculation

For each set of the human SZFP gene orthologs, multiple sequence alignments were constructed using the codon_alignment.pl script (*82*) (https://github.com/lyy005/codon_alignment/blob/master/) and used as input for the HyPhy pipeline. Specifically, we first generated phylogenetic trees of the orthologs using PhyML (*73*) (v3.3.20220408). These trees were then analysed with the HyPhy (*83*) (version 2.5.2) pipeline, utilising the FEL subroutine to calculate the *d*_N_/*d*_S_ ratios and to determine their significance for each nucleotide position in the SZFP genes with a *P* value cutoff of 0.05.

#### MNase-seq analysis

Raw read mapping to hg38 was performed with Bowtie2 (*75*) (version 2.5.1), with the “--very-sensitive-local --no-discordant” options. The resulting alignment BAM files were used as input for the dpos function from DANPOS3 (*84*) (version 3.0.0) to obtain genome-wide nucleosome occupancy values and to detect nucleosomal positions for each condition. For the heatmap representation of occupancy around specific sets of genomic loci, we used the plotHeatmap tool from deepTools (*85*) (version 3.5.5).

#### ATAC-seq analysis

Raw read mapping to hg38 or mm10 and peak calling were performed with PEPATAC (*86*) with the default parameters. The ATAC-seq peaks called in all the samples were merged with the bedtools (*87*) (v2.30.0) merge function, and reads overlapping to these loci were counted with bedtools (v2.30.0) multicov function. For the accessibility analysis of TEs, TE annotation obtained from UCSC genome browser was used instead of ATAC-seq peaks for signal quantification. The resulting read counts were used as an input for differential accessibility analysis with DESeq2 (*88*) (version 1.34.0). For the genome browser visualisation of the data, we used plotgardener (*89*) (version 1.0.17), and, for the heatmap representation of signals around specific sets of genomic loci, we used the plotHeatmap tool from deepTools (*85*) (version 3.5.5).

#### RNA-seq analysis

Raw read mapping to hg38 or mm10, filtering, and per 3′ UTR read quantification were performed with SLAM-DUNK (*90*) (version 0.4.3) with the parameters “-5 12 -n 100 -m -rl 75 -q --skip-sam”. The resulting read counts were used as an input for differential gene expression analysis with DESeq2 (*88*) (version 1.34.0).

## Acknowledgements

We thank the Trono lab members for constructive discussions. Most computational works were conducted on the high-performance computing cluster developed and maintained by Scientific IT and Application Support (SCITAS) at EPFL. We thank the Gene Expression Core Facility (GECF) and Flow Cytometry Core Facility (FCCF) at EPFL for technical support.

## Funding

European Research Council No. 268721 (DT)

European Research Council No. 694658 (DT)

Swiss National Science Foundation 310030_152879 (DT)

Swiss National Science Foundation 310030B_173337 (DT)

European Molecular Biology Organization (EMBO) Postdoctoral Fellowship

ALTF1287-2020 (WM)

Japan Society for the Promotion of Science (JSPS) Overseas Research Fellowship no. 202360326 (WM)

## Author contributions

Conceptualization: WM, DT

Data curation: WM

Formal analysis: WM, JD, SS, CP, DG, EP

Funding acquisition: WM, DT

Investigation: WM, SO, CR

Methodology: WM, JD, CP

Project administration: WM, DT

Resources: EP

Software: WM, JD, SS, CP

Supervision: WM, EP, DT

Validation: WM

Visualization: WM

Writing – original draft: WM, JD

Writing – review & editing: WM, DT, CP, JD

## Data and materials availability

High throughput sequencing data were deposited on ArrayExpress (https://www.ebi.ac.uk/biostudies/arrayexpress) under the following accession numbers: SZFP ChIP-seq (E-MTAB-14683), *ZNF24*^*FKBP*^ MNase-seq (E-MTAB-14880), *ZNF24*^*FKBP*^ ATAC-seq (E-MTAB-14712), *ZNF24*^*FKBP*^ ChIP-mentation (E-MTAB-14699), *ZNF24*^*FKBP*^ RNA-seq (E-MTAB-14711), *Zscan10*^*FKBP*^ ATAC-seq (E-MTAB-14702), *Zscan10*^*FKBP*^ ChIPmentation (E-MTAB-14689), *Zscan10*^*FKBP*^ RNA-seq (E-MTAB-14692).

---

**Figure S1:**
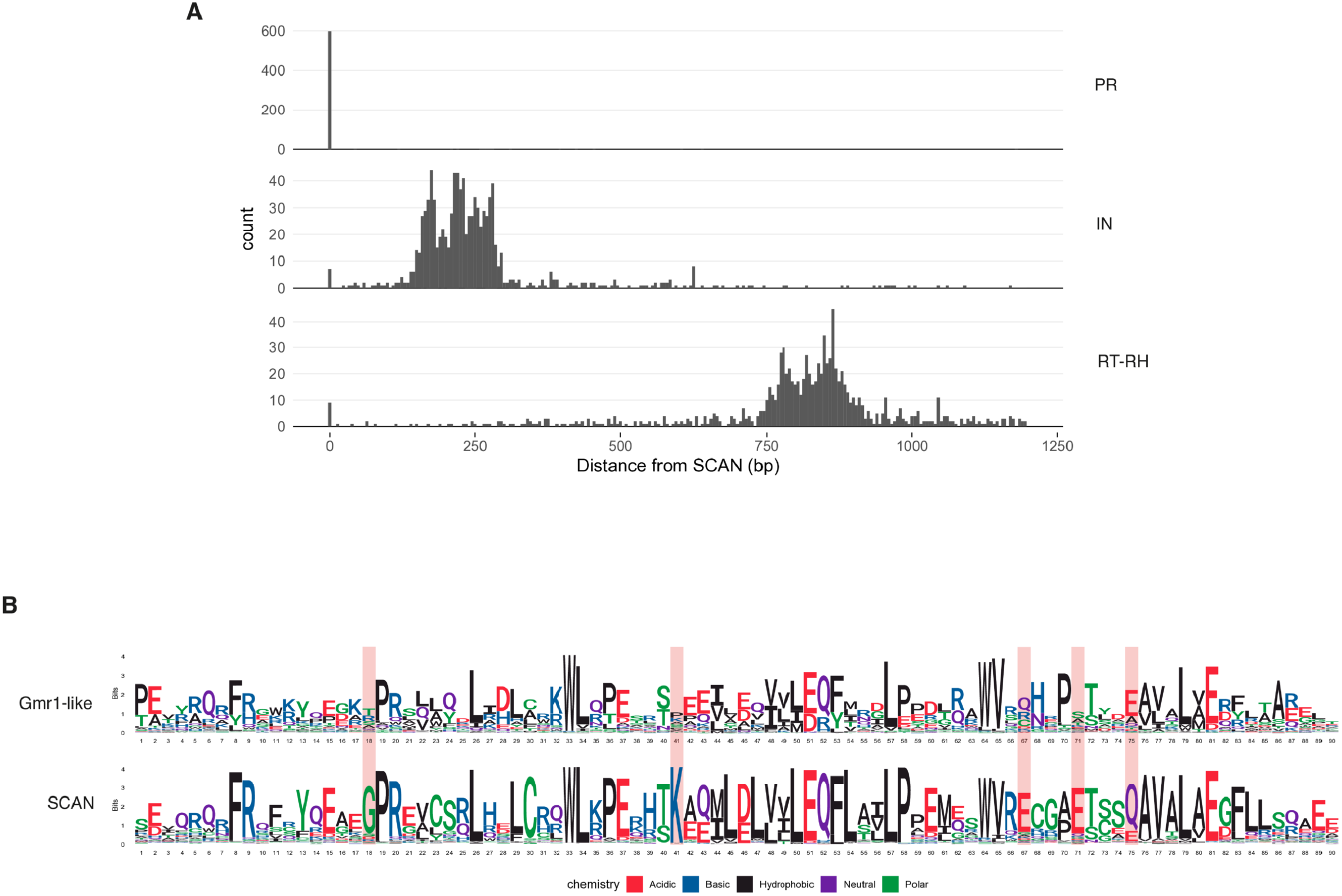
Characterisation of SCAN-encoding elements in vertebrates. (A) Histograms summarising distances to different retroviral domains found downstream of SCAN domains. (B) Amino-acid sequence motifs of the SCAN domains encoded by Gmr-like elements and ZF-associated genes in the representative vertebrate genomes used in Fig. 1C. Red shading highlights residues that differ between the two.

**Figure S2:**
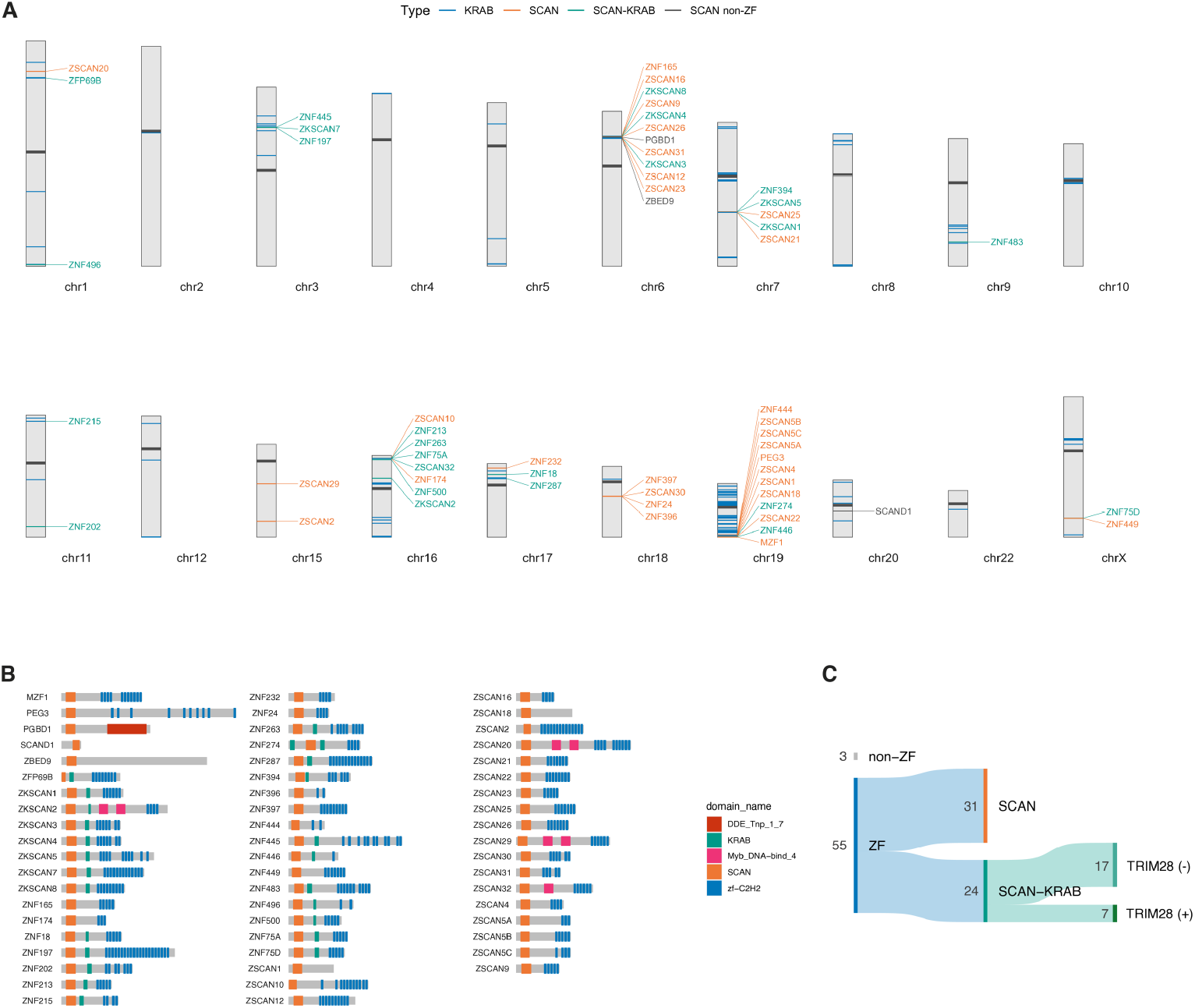
Genetic map and domain organisation of the human SZFPs. (A) Idiogram of the human SCAN-encoding genes (labelled with their gene name) and KZFP genes. Only chromosomes encoding these genes are shown. (B) Protein domain organisation of the human SCAN-containing proteins. (C) A Sankey diagram summarising the different domain compositions of the human SCAN-containing proteins. SZFPs with a KRAB domain are further grouped based on their TRIM28 interaction.

**Figure S3:**
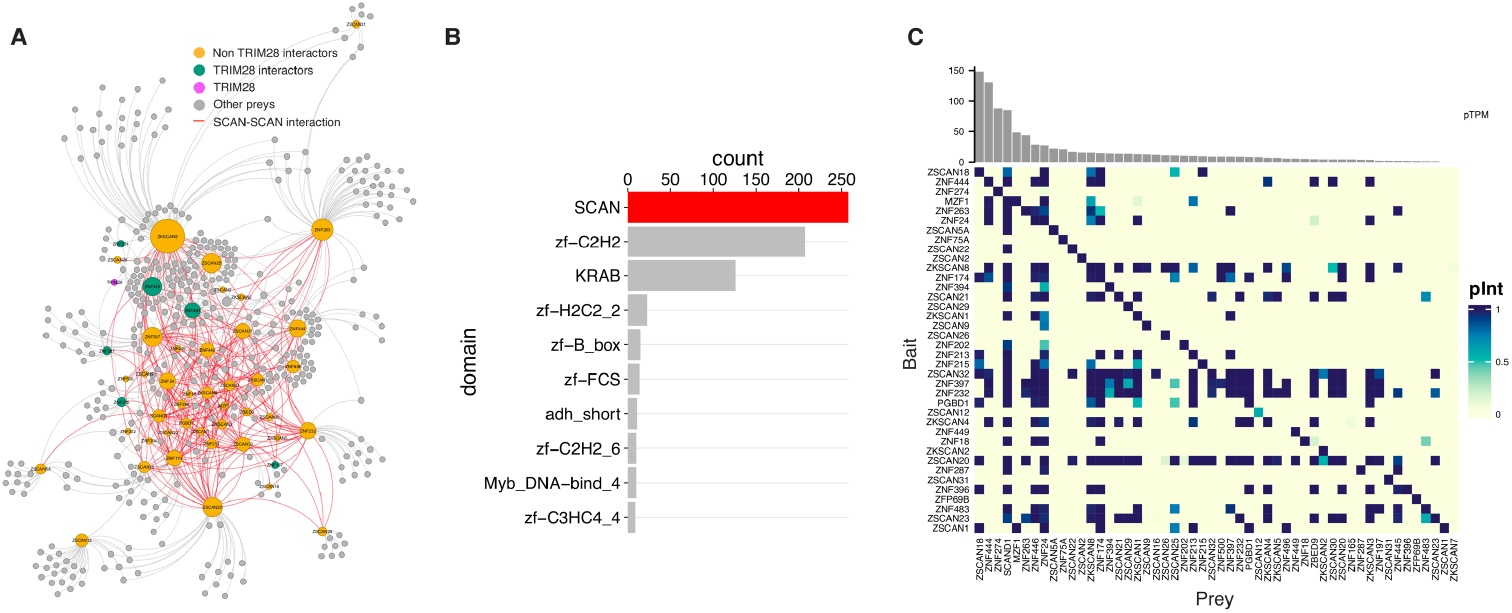
Protein interactome of the human SCAN-containing proteins. (A) A protein-protein interaction network of the human SCAN-containing proteins based on Bioplex 3.0. Significant interactions are shown as lines linking nodes, which correspond to proteins. Only SCAN-containing proteins and TRIM28 are labelled with their protein name. (B) Frequency of domains annotated in interactors of SCAN-containing proteins. (C) A heatmap summarising the protein-protein interaction probability (pInt) of SCAN domain-containing proteins obtained from Bioplex 3.0. The expression levels of the prey proteins in 293T are shown at the top.

**Figure S4.**
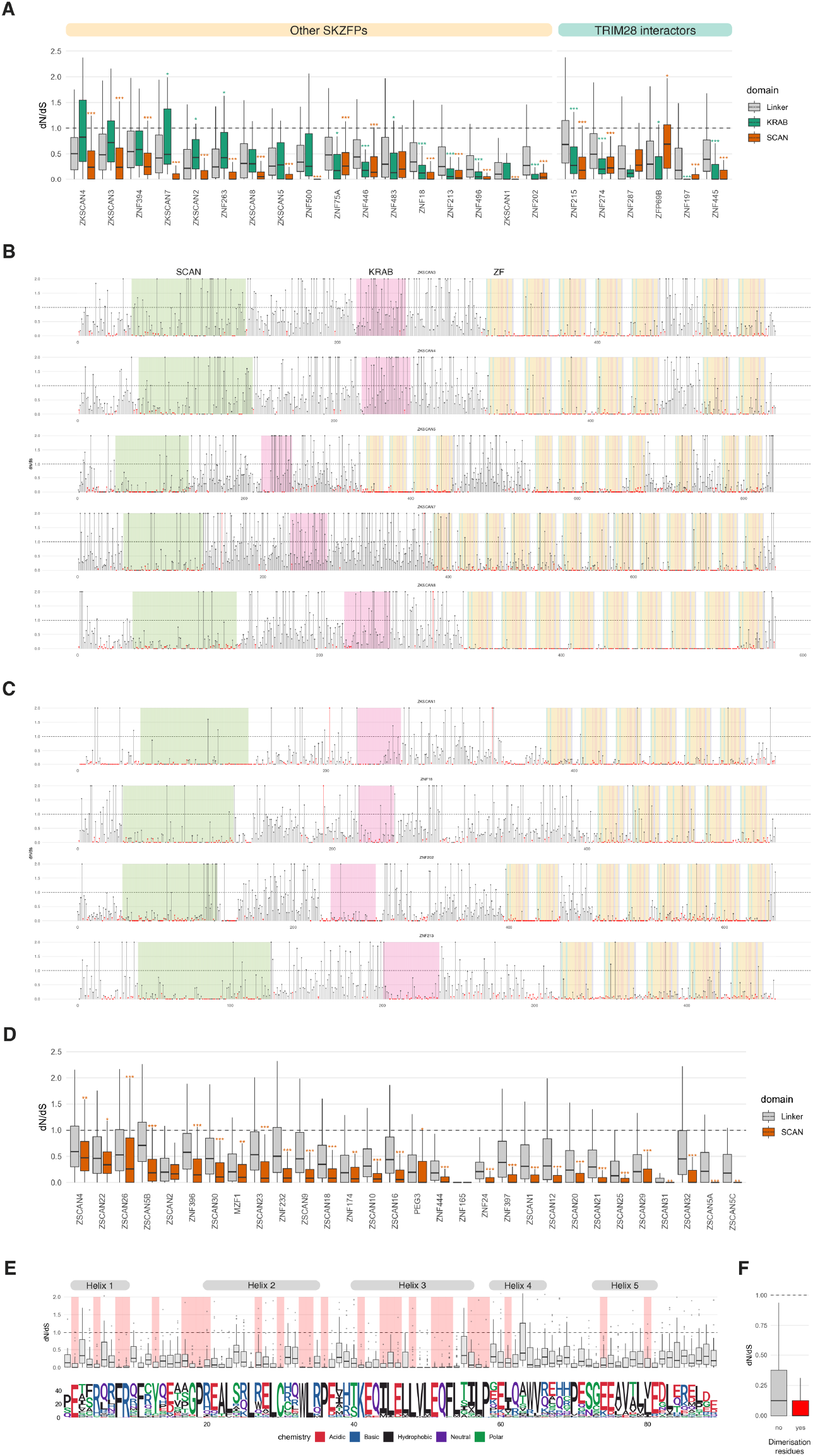
Differential selection on amino-acid residues in the human SZFPs. (A) Boxplots of amino-acid level *d*_N_/*d*_S_ values in distinct domains of human ZFPs with both SCAN and KRAB domains. “Linker” refers to protein segments where none of the following is annotated: SCAN, KRAB, and ZF domains. Mann-Whitney *U* tests were performed comparing the linker region to either the KRAB or SCAN domain and significant *P* values are indicated as following: *, *P* < 0.05; **, *P* < 0.01; ***, *P* < 0.001. (B, C) *d*_N_/*d*_S_ values over amino-acid residues of SZFPs containing a KRAB domain. Those containing a KRAB with few residues under purifying selection (B) and with a number of residues under significant purifying selection (C). Residues under significant positive or negative selection are shown in red. Coloured shades highlight protein domains. (D) Boxplots showing the *d*_N_/*d*_S_ distributions over the linker- and the SCAN-encoding regions of KRAB-less SZFP genes. Mann-Whitney *U* tests were performed comparing the linker region to either the KRAB or SCAN domain, and significant *P* values are shown as following: *, *P* < 0.05; **, *P* < 0.01; ***, *P* < 0.001. (E) Boxplots of *d*_N_/*d*_S_ distribution (top) and the frequency of amino acids in the human SZFPs (bottom) for each SCAN residue. Grey bars indicate residues forming helices of SCAN, and red shades highlight those that are involved in dimerisation. (F) Boxplot comparing *d*_N_/*d*_S_ of the SCAN residues grouped based on their involvement in dimerisation.

**Figure S5.**
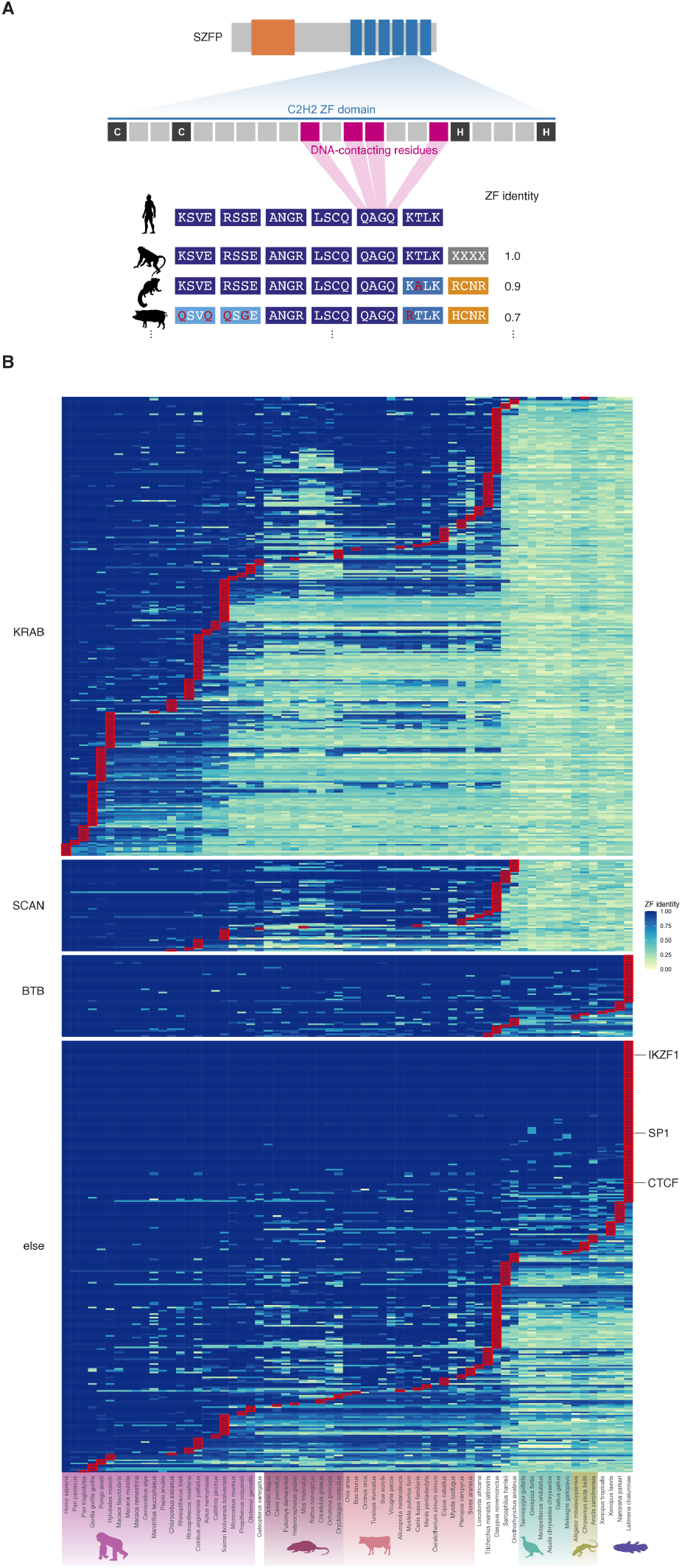
Evolutionary trajectory of the DNA-contacting residues of the human ZFPs. (A) A schematic representation of the pairwise ZF comparison. For each human ZFP, DNA-contacting residues (ZFprint) were compared against those in another genome. A ZF identity score was calculated by dividing the number of matched residues by the total number. (B) A heatmap summarising ZF identity scores of the human SZFPs compared to those encoded in other vertebrate genomes. Red borders highlight the most distant species that harbours a ZFP with an identical ZFprint.

**Figure S6.**
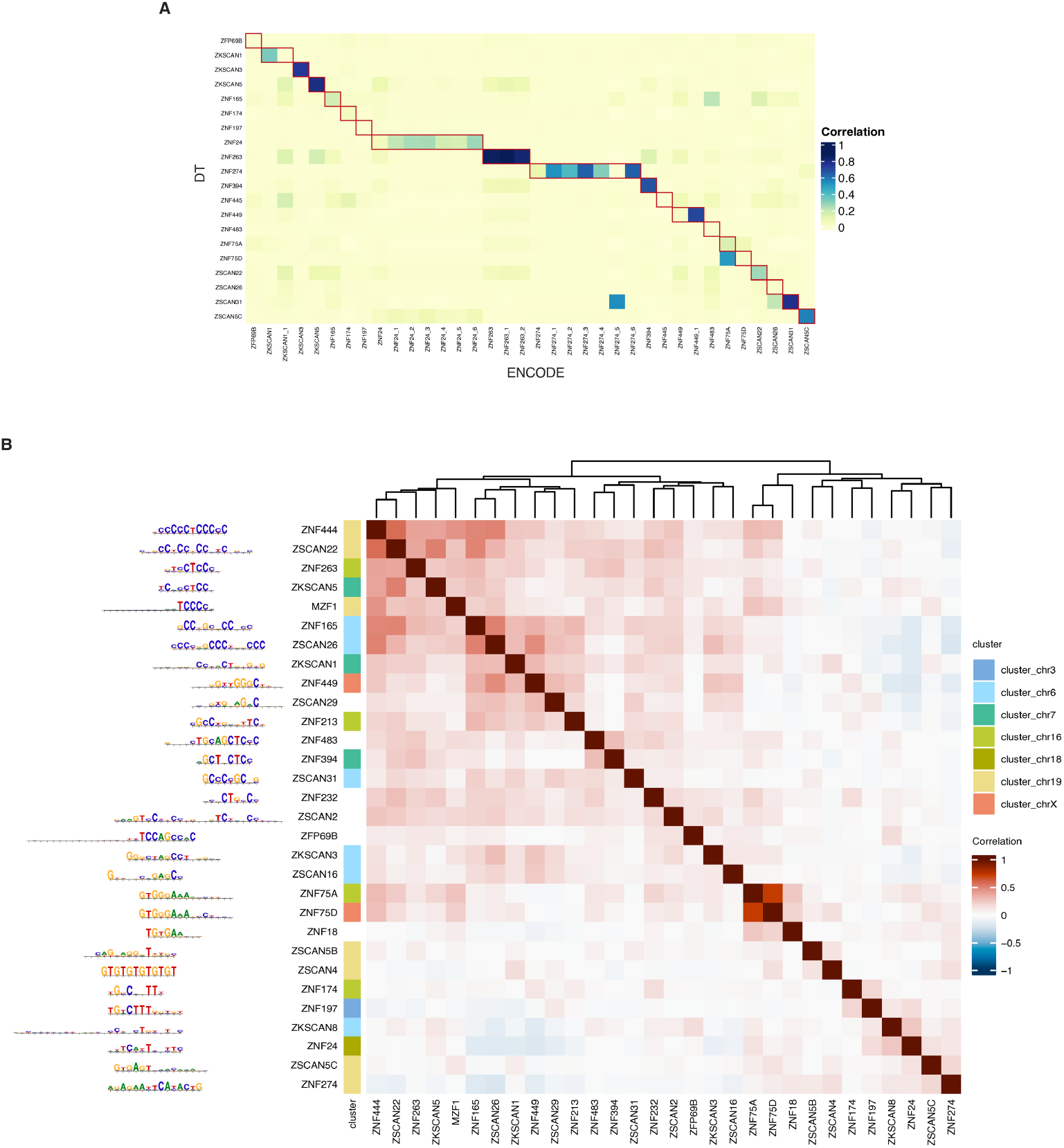
SZFP binding motifs determined by ChIP-seq. (A) Heatmap summarising similarity between SZFP motifs identified in this study (DT) and those by ENCODE. The similarity scores were calculated using MoSBAT. (B) Pairwise motif similarity of the determined human SZFP motifs. Genes encoded as part of an SZFP gene cluster are labelled with the cluster name (see also fig. S2A).

**Figure S7.**
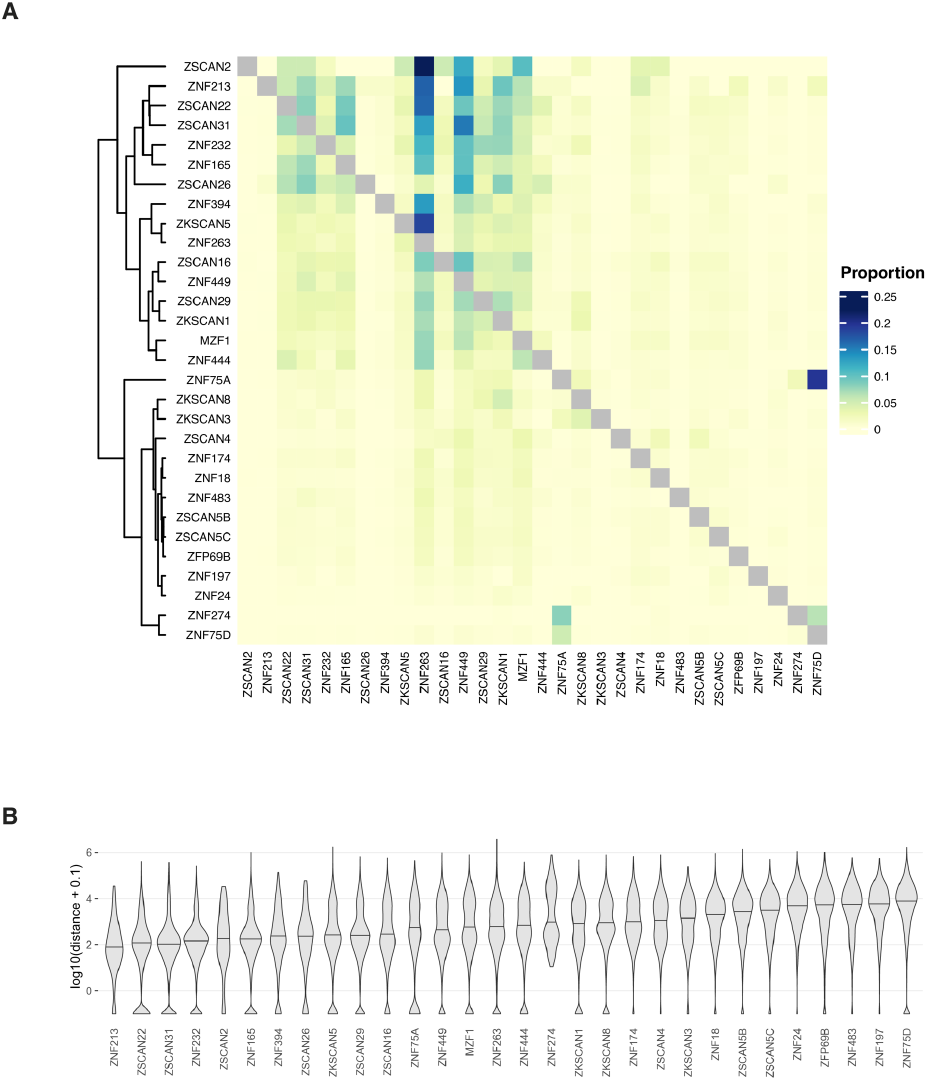
Genome-wide distribution of the SZFP binding sites. (A) Heatmap summarising similarity between SZFP motifs identified in this study (DT) and those by ENCODE. The similarity scores were calculated using MoSBAT. (B) Pairwise motif similarity of the determined human SZFP motifs. Genes encoded as part of an SZFP gene cluster are labelled with the cluster name (see also fig. S2A).

**Figure S8.**
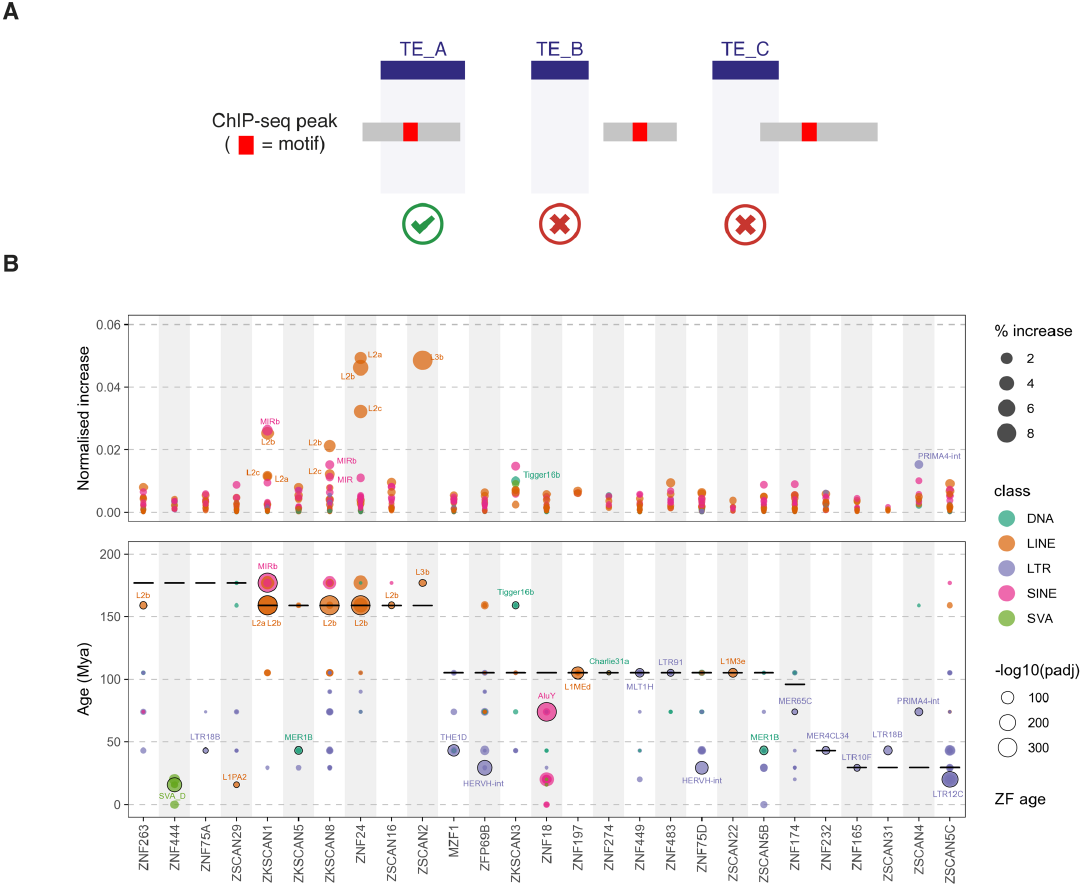
Recognition of TEs and degTEs by human SZFPs. (A) A schematic of the pipeline of calling overlap between SZFP ChIP-seq peaks and TEs. An overlap is called only when a binding motif within a ChIP-seq peak overlaps a TE integrant. (B) Increase in TE-overlapping peaks after the addition of degTEs normalised to the number of peaks (top) shown with significantly overlapping TE subfamilies (bottom).

**Figure S9.**
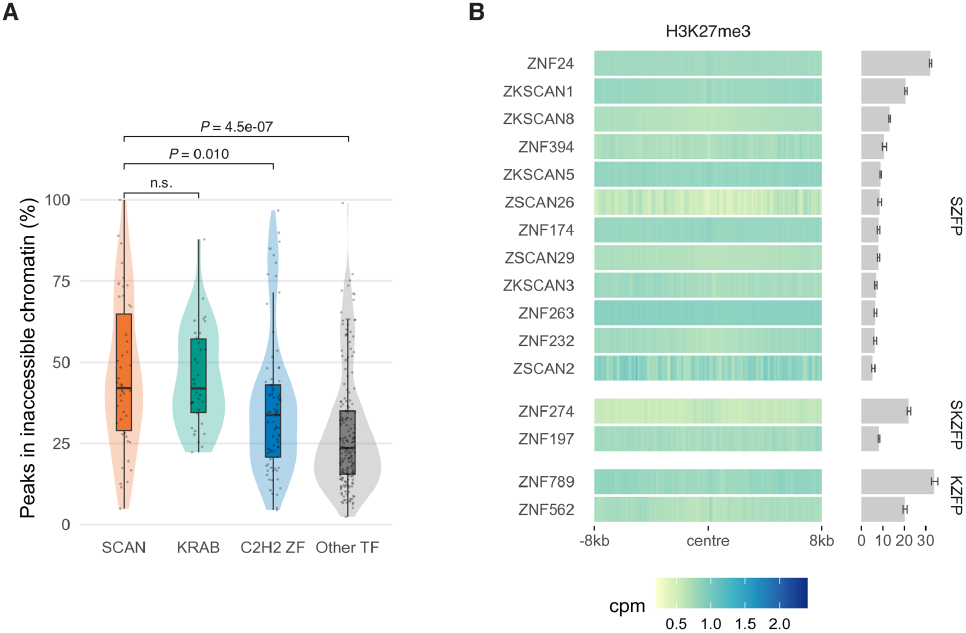
Epigenetic profiles of the SZFP binding sites compared to other TFs. (A) Violin plots of proportions of peaks in inaccessible chromatin determined by ENCODE ChIP-seq and ATAC-seq performed in K562. TFs were grouped according to their domain configuration: SCAN, any TFs with a SCAN domain; KRAB, TFs with a KRAB domain and without a SCAN domain; C2H2 ZF, ZFPs with neither SCAN nor KRAB domain. *P* values of a Mann-Whitney *U* test are shown. (B) H3K27me3 signals around the binding sites of ZFPs expressed in K562.

**Figure S10.**
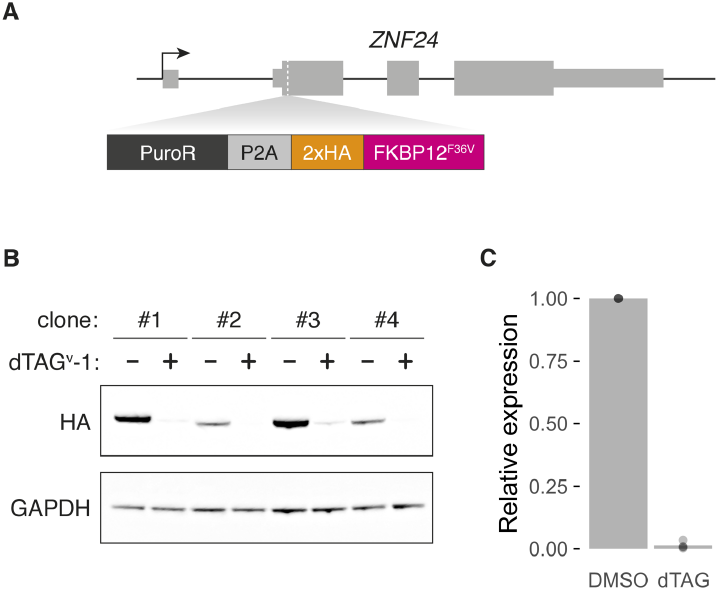
Establishment of the *ZNF24*^*FKBP*^ HAP1 cell lines. (A) Schematic representation of the *ZNF24* gene tagging with FKBP12^F36V^. (B) Western blot analysis of protein levels of endogenously-tagged ZNF24 and GAPDH in four clones of *ZNF24*^*FKBP*^ cell lines with or without exposure to dTAG^v^-1, using anti-HA antibody and anti-GAPDH antibody, respectively. (C) Quantitative comparison of the ZNF24 protein levels normalised by GAPDH based on the Western blot signal shown in (B). For the image analysis of the blot, we used GelAnalyzer 23.1.1 (available at www.gelanalyzer.com) by Istvan Lazar Jr., PhD and Istvan Lazar Sr., PhD, CSc.

**Figure S11.**
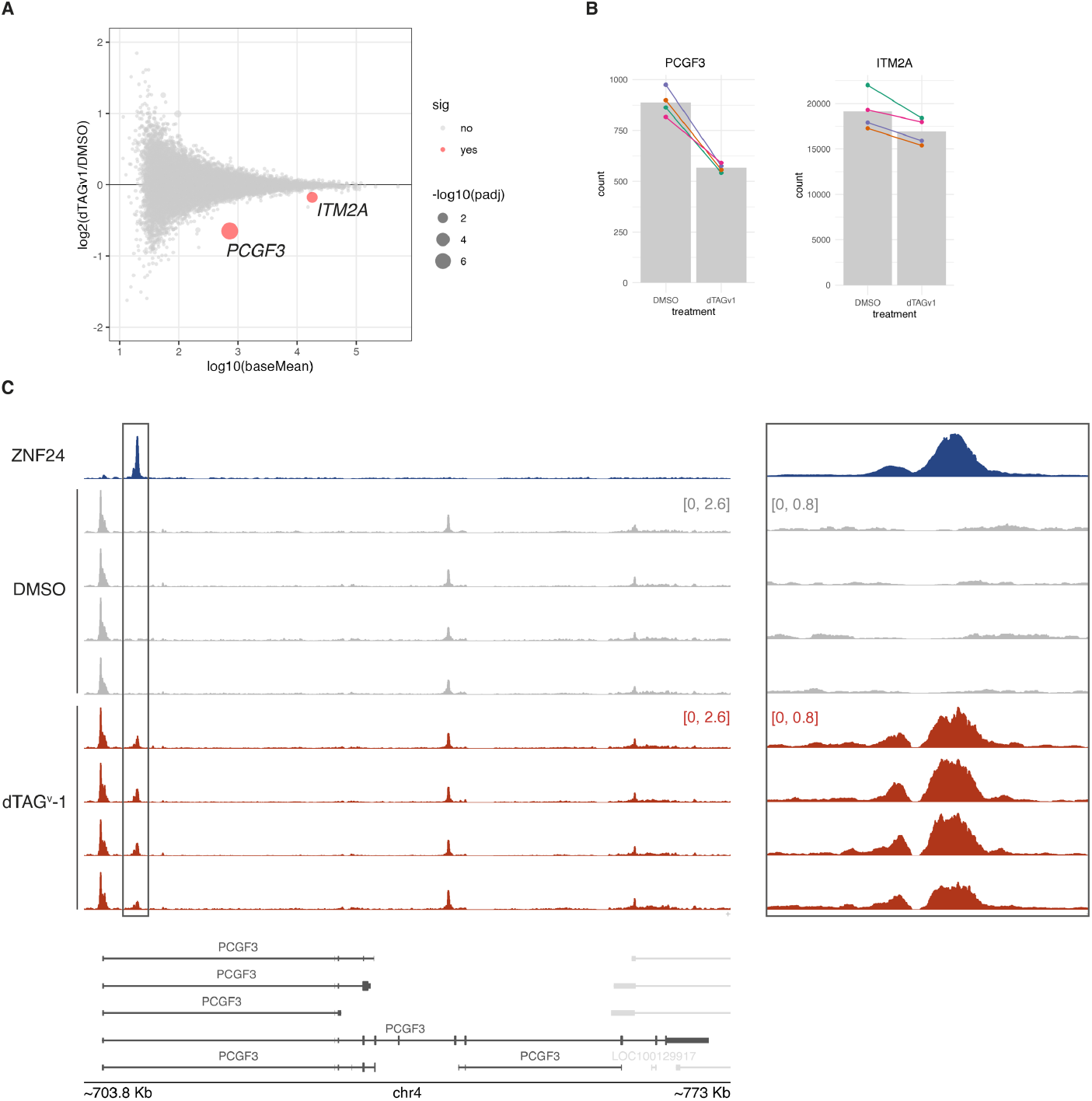
Effects of ZNF24 depletion on transcriptome. (A) MA plot summarising the transcriptional changes upon ZFN24 depletion measured by RNA-seq. (B) DESeq2-normalised read counts of the two differentially expressed genes. Colours represent different *ZNF24*^*FKBP*^ clones. (C) ZNF24 ChIP-seq and ATAC-seq profiles (CPM) around the *PCGF3* locus comparing the cells with and without ZNF24 depletion by dTAG^v^-1. Genes shown in light grey are encoded on the reverse strand.

**Figure S12.**
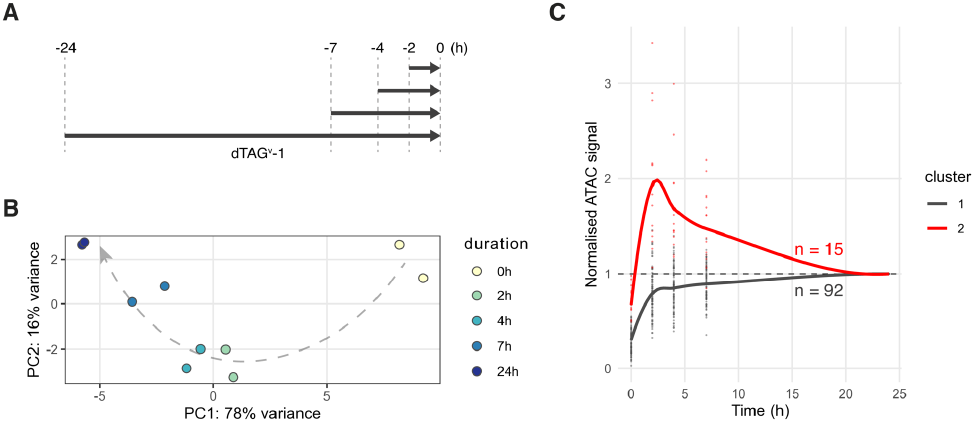
Time-course chromatin accessibility changes induced by ZNF24 depletion. (A) Scheme summarising the design of the time-course experiment. *ZNF24*^*FKBP*^ cell lines exposed to dTAG^v^-1 for different durations were harvested at the same time for ATAC-seq library preparation. (B) The first two principal components (PCs) of the normalised read counts of the samples with different dTAG^v^-1 exposure times. (C) Relative ATAC-seq signals around the opened regions normalised to the 24-hour time point. The regions were grouped into two clusters based on k-means clustering.

**Figure S13.**
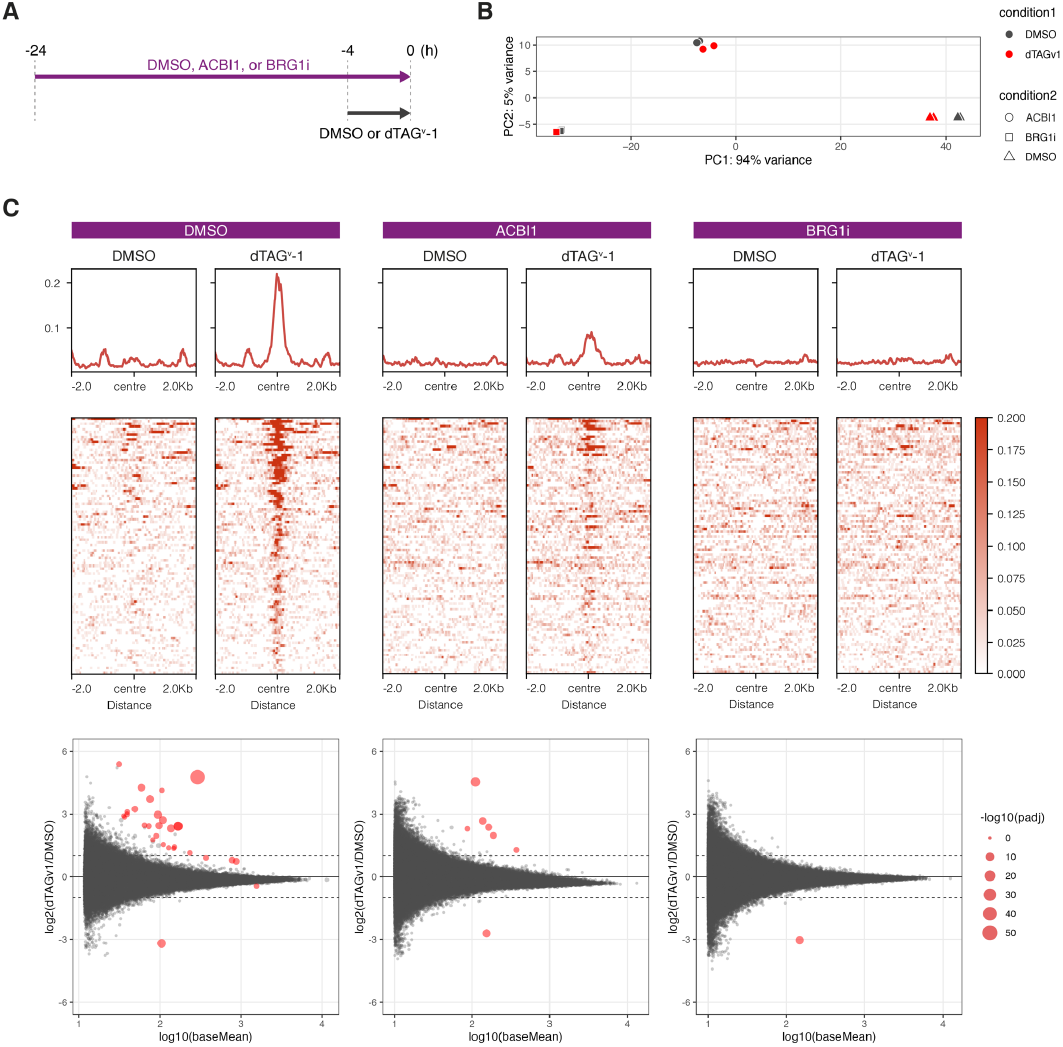
SWI/SNF dependency of chromatin accessibility changes induced by ZNF24 depletion. (A) Scheme summarising the design of the SMARCA2/4 perturbation experiment. *ZNF24*^*FKBP*^ cell lines were treated with different drugs that perturb SMARCA2/4 prior to exposure to dTAG^v^-1 or DMSO. (B) The first two principal components (PCs) of the normalised read counts of the samples treated with different SMARCA2/4-perturbing agents. (C) Average profiles of ATAC-seq signals (CPM) around the opened regions (top) and MA plots summarising the ATAC-seq signal changes upon ZNF24 depletion (bottom).

**Figure S14.**
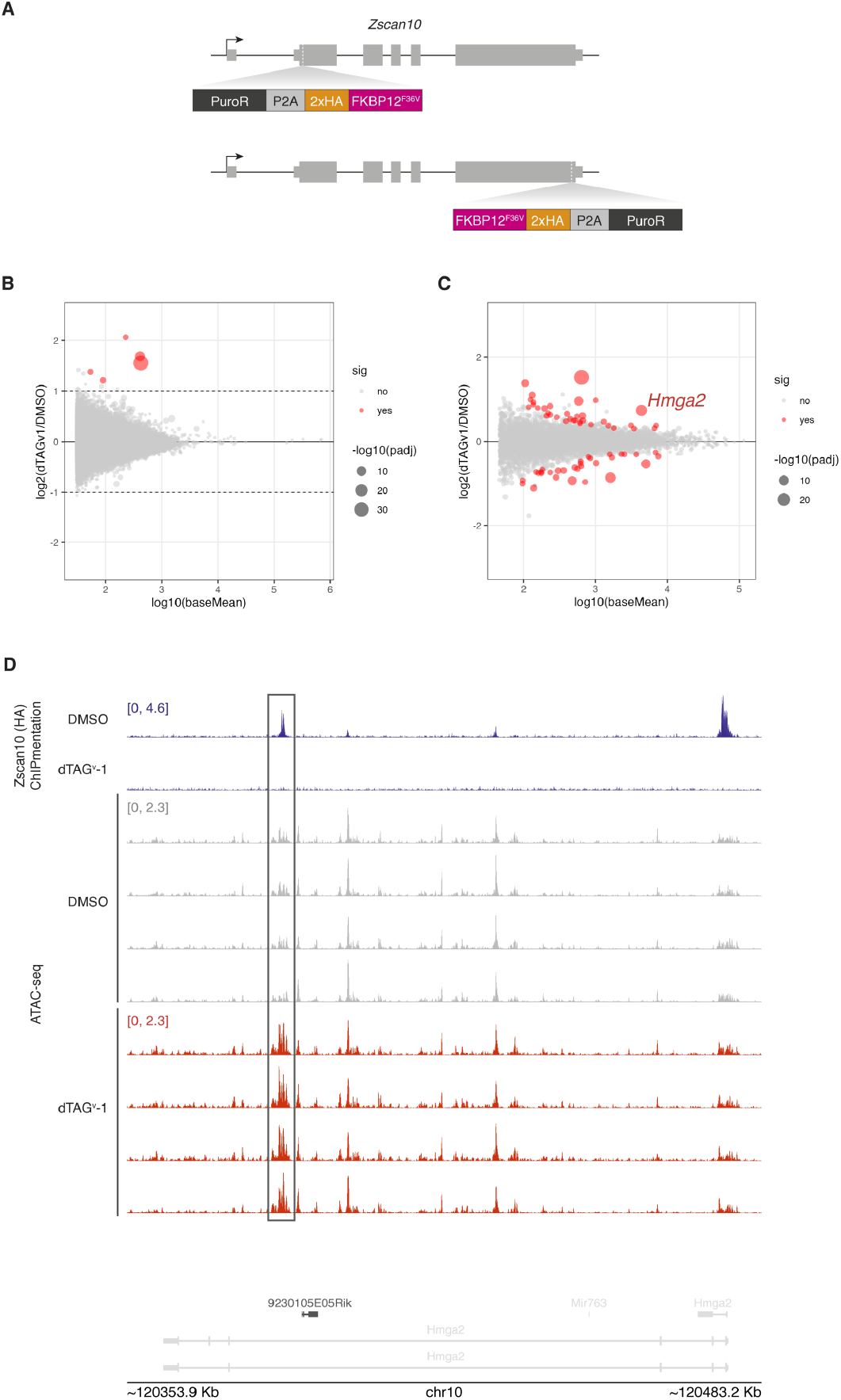
Transcriptomic and chromatin accessibility changes upon Zscan10 depletion in mESCs. (A) Endogenous tagging strategies of the mouse *Zscan10* gene with an FKBP12^F36V^ degron. Two clones of each N- and C-terminally tagged mESC lines were established. (B) MA plot summarising the ATAC-seq change upon Zscan10 depletion. (C) MA plot summarising the RNA-seq change upon ZSCAN10 depletion. (D) ZSCAN10 ChIPmentation and ATAC-seq signal (CPM) changes around the *Hmga2* locus. Genes shown in light grey are encoded on the reverse strand.

**Table S1.** Meta-analysis of SZFP-TRIM28 interaction.

| gene | AA | Itokawa | Tycko | Helleboid | Bioplex | TRIM28 interaction call (this study) |
| --- | --- | --- | --- | --- | --- | --- |
| ZFP69B | s | NA | TRUE | NA | FALSE | TRUE |
| ZKSCAN1 | v | FALSE | FALSE | NA | FALSE | FALSE |
| ZKSCAN2 | v | NA | FALSE | NA | FALSE | FALSE |
| ZKSCAN3 | v | NA | FALSE | FALSE | FALSE | FALSE |
| ZKSCAN4 | v | NA | FALSE | FALSE | FALSE | FALSE |
| ZKSCAN5 | v | NA | FALSE | FALSE | FALSE | FALSE |
| ZKSCAN7 | v | NA | FALSE | FALSE | NA | FALSE |
| ZKSCAN8 | v | NA | FALSE | FALSE | FALSE | FALSE |
| ZNF18 | v | FALSE | FALSE | FALSE | FALSE | FALSE |
| ZNF197 | s | NA | TRUE | TRUE | NA | TRUE |
| ZNF202 | v | FALSE | NA | FALSE | FALSE | FALSE |
| ZNF213 | v | FALSE | FALSE | FALSE | FALSE | FALSE |
| ZNF215 | s | NA | NA | TRUE | FALSE | TRUE |
| ZNF263 | v | NA | FALSE | FALSE | FALSE | FALSE |
| ZNF274 | s | NA | TRUE | TRUE | TRUE | TRUE |
| ZNF287 | s | TRUE | NA | TRUE | TRUE | TRUE |
| ZNF394 | v | NA | FALSE | NA | FALSE | FALSE |
| ZNF445 | s | TRUE | TRUE | TRUE | TRUE | TRUE |
| ZNF446 | v | FALSE | FALSE | FALSE | NA | FALSE |
| ZNF483 | s | NA | NA | NA | FALSE | FALSE |
| ZNF496 | v | NA | FALSE | FALSE | FALSE | FALSE |
| ZNF500 | v | FALSE | FALSE | NA | NA | FALSE |
| ZNF75A | v | NA | NA | NA | FALSE | FALSE |
| ZNF75D | v | NA | TRUE | TRUE | NA | TRUE |

**Table S2.** Data source of SZFP binding sites.

| gene | source | cell line | accession |
| --- | --- | --- | --- |
| MZF1 | ENCODE | 293T | GSE127576 |
| ZFP69B | DT | 293T |  |
| ZKSCAN1 | ENCODE | K562 | GSE92129 |
| ZKSCAN3 | ENCODE | K562 | GSE170739 |
| ZKSCAN5 | ENCODE | HepG2 | GSE170522 |
| ZKSCAN8 | ENCODE | K562 | GSE105606 |
| ZNF165 | DT | 293T |  |
| ZNF174 | DT | 293T |  |
| ZNF18 | ENCODE | 293T | GSE92202 |
| ZNF232 | ENCODE | K562 | GSE231043 |
| ZNF24 | ENCODE | GM12878 | GSE105225 |
| ZNF263 | ENCODE | HepG2 | GSE170334 |
| ZNF274 | ENCODE | HCT116 | GSE136460 |
| ZNF394 | DT | 293T |  |
| ZNF444 | DT | 293T |  |
| ZNF449 | DT | 293T |  |
| ZNF483 | DT | 293T |  |
| ZNF75A | ENCODE | K562 | GSE174983 |
| ZNF75D | DT | 293T |  |
| ZSCAN16 | ENCODE | 293T | GSE92115 |
| ZSCAN2 | DT | 293T |  |
| ZSCAN22 | DT | 293T |  |
| ZSCAN26 | DT | 293T |  |
| ZSCAN29 | ENCODE | K562 | GSE91549 |
| ZSCAN31 | DT | 293T |  |
| ZSCAN4 | DT | 293T |  |
| ZSCAN5B | DT | 293T |  |
| ZSCAN5C | DT | 293T |  |

